# Sex-specific *doublesex* regulation targeting the color-patterning gene *h* underlies the evolution of wing sexual dimorphism in the harlequin ladybug *Harmonia axyridis*

**DOI:** 10.1101/2025.08.06.668842

**Authors:** Soichi Yeki, Kagayaki Kato, Shinichi Morita, Kenji Shimomura, Teruyuki Niimi, Norihide Hinomoto, Takaaki Daimon, Toshiya Ando

**Author notes:** Correspondence and requests for materials should be addressed to T.A.

## Abstract

Organisms on Earth show various forms of sexual dimorphism, including ornaments, weapon traits, and pheromone glands, which has been acquired through sexual selection during evolution. Although the genetic basis of sexual trait has been investigated in diverse species, its evolutionary origin, evolutionary mode, and evolvability remain poorly understood. To address these issues, we investigated the strain-specific sexual dimorphism in elytral color patterns of the harlequin ladybug, *Harmonia axyridis*, a species with over 200 color morphs. The most basal Red-nSpots type color morph exhibits sexual dimorphism, whereas other derived color morphs have lost it. To investigate how this loss of sexual dimorphism is associated with the emergence of novel color morphs, we investigated the genetic basis of sexual dimorphism by focusing on the master sex differentiation gene, *doublesex* (*dsx*). We show that *dsx* regulates color pattern dimorphism by negatively modulating black spot size in males. This modulation is primarily mediated by the transcriptional regulation of the color patterning gene, *h* (*Drosophila pannier* ortholog). Intraspecific comparative ATAC-seq analysis of the pupal wings revealed that, at the *h* locus, not the absolute number of Dsx-binding motifs but the proportion of open chromatin regions containing Dsx-binding motifs relative to those lacking such motifs was reduced in strains that had lost sexual dimorphism and acquired novel color patterns, implying that sexual dimorphism evolves based on the balance between novel CREs and Dsx-binding motif density. The present study provides a fundamental molecular framework for understanding how a secondary sexual trait evolves within *H. axyridis*.

## Introduction

Organisms on Earth exhibit a wide range of secondary sexual traits, including ornaments, weapon traits, and pheromone glands. These traits have evolved in addition to primary reproductive structures (e.g., gonads that produce gametes) and contribute to reproductive fitness in diverse environments. Recent studies on the molecular basis of secondary sexual trait formation in various insects have revealed both conserved and divergent mechanisms (Hopkins & Kopp, 2021). In most insects, the conserved sex differentiation gene *doublesex* (*dsx*), which encodes a DMRT (Doublesex and Mab-3-Related Transcription factor) type transcription factor (Burtis & Baker, 1989; Raymond et al., 1998), regulates a wide array of sexual traits ranging from primary sexual structures to secondary sexual traits such as ornaments. However, the regulatory mode, timing of action, and downstream targets of *dsx* in secondary sexual trait formation vary significantly across species (Verhulst & van de Zande, 2015). These findings suggest that the ancestral *dsx*-mediated sex differentiation pathway has been repeatedly recruited (co-opted) to facilitate the evolution of secondary sexual traits in divergent insect lineages. The molecular mechanisms underlying this regulatory co-option have been actively studied in *Drosophila* sister species, focusing on species-specific secondary sexual traits, such as abdominal pigmentation (Kopp, Duncan, & Carroll, 2000; Williams et al., 2008), sex combs on the forelegs (Tanaka, Barmina, Sanders, Arbeitman, & Kopp, 2011), and cuticular carbohydrate deposition (Shirangi, Dufour, Williams, & Carroll, 2009). Still, the diverse evolutionary strategies for sexual traits in organisms highlight the need for comparative analyses in other insect lineages to understand the universal principles of sexual trait evolution.

To address this issue, we focused on the sexual dimorphism of the forewing (elytral) color pattern in the harlequin ladybug, *Harmonia axyridis*. *H. axyridis* displays over 200 intraspecific color morphs, which are believed to be regulated by a combination of more than 22 alleles at a single genetic locus, *h* (reviewed in (Komai, Chino, & Hosino, 1950)). Of those alleles, the most recessive class, *h^succinea^* (collectively abbreviated as *h*), exhibits sexual dimorphism. The *h* alleles exhibit zero to ten black spots on the red background in the single forewing (Fig. 1A, hereafter Red-nSpots). Females have larger black spots than males (Hosino, 1942, 1948; Knapp & Nedvěd, 2013) (Fig. 1B, 2-10). In contrast, other major *h* alleles (*h^Conspicu^* [*h^C^*], *h^Spectabilis^* [*h^Sp^*], and *h^Axyxirids^* [*h^A^*]), which show one, two, and six red spots on the black background (Fig. 1A, hereafter Black-2Spots, Black-4Spots, and Black-nSpots [6–12 spots in the wild], respectively), do not exhibit conspicuous sexual dimorphism. This allele-specific sexual dimorphism within a species presents an excellent opportunity to investigate the evolutionary dynamics of secondary sexual traits. Previous genomic studies have identified *h* as the ortholog of the *Drosophila pannier* gene, which encodes a GATA transcription factor, and revealed that the non-coding sequences in its first intron are highly divergent within a species (Ando et al., 2018; Gautier et al., 2018). In the present study, we refer to the *Harmonia* ortholog of the *Drosophila pannier* gene as *h*. The molecular phylogenetic analysis using the conserved non-coding sequences in the *h* intron suggested that the Red-nSpots allele is the basal allele, and that the other alleles (Black-2Spots, Black-4Spots, and Black-nSpots) are more recently diverged (Ando et al., 2018) (Fig. 1A). Moreover, the Red-nSpots-like patterns prevail in the genus *Harmonia* (e.g., *H. octomaculata*, *Harmonia conformis*, *Harmonia quadripunctata*), implying that the *h* allele retains ancestral-like features and that the sexual dimorphism was somehow lost during evolution. Therefore, elucidating the genetic basis of sexual dimorphism in the Red-nSpots allele provides a valuable framework to explore the origin, mechanisms, and evolvability of secondary sexual traits in *H. axyridis*.

**Fig. 1.**
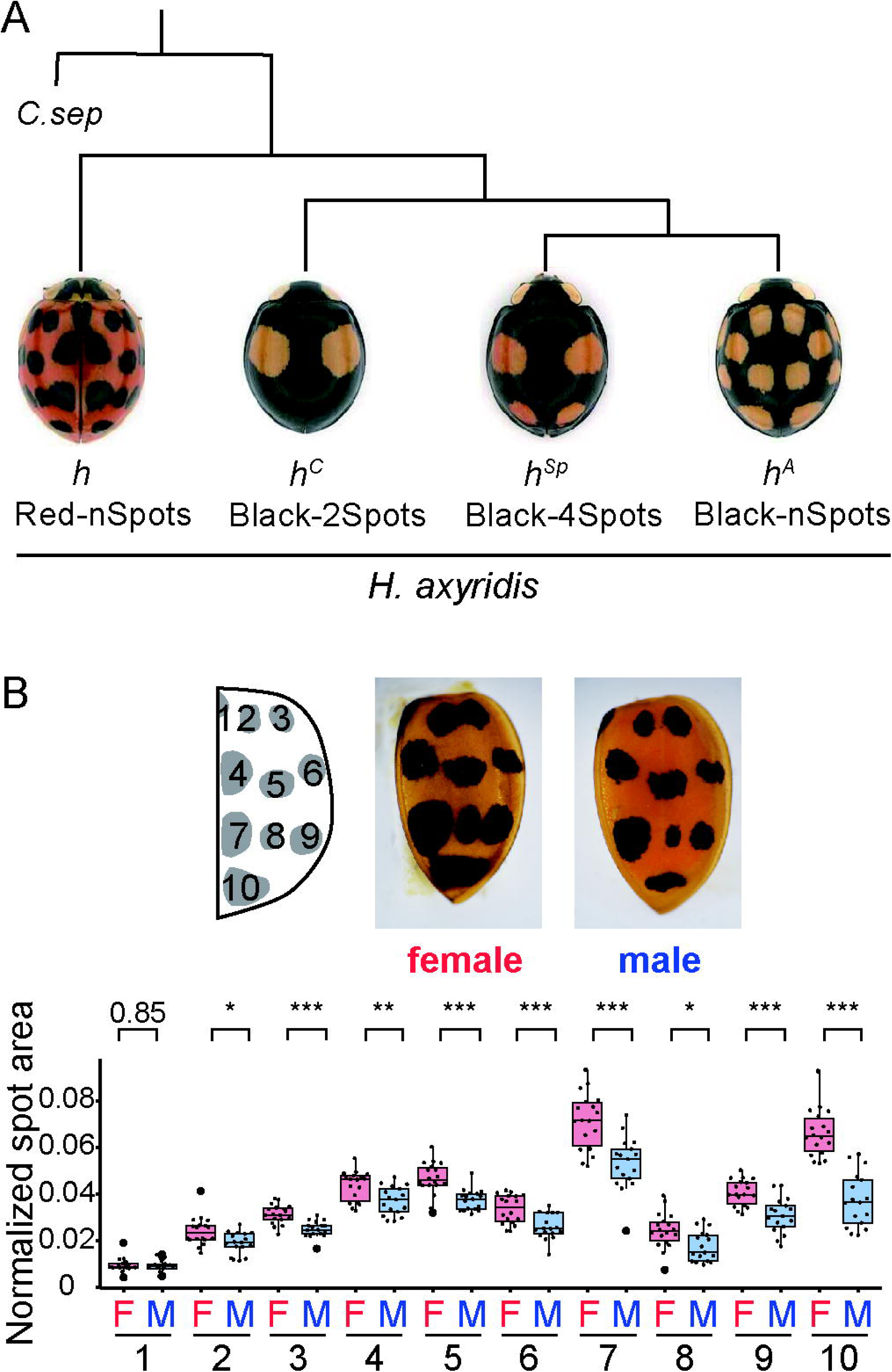
Intraspecific genetic polymorphism of elytral color patterns in *H. axyridis.* **(A)** Representative phenotypes and phylogenetic relationships of four major *h* alleles responsible for elytral color pattern variation (*h*, succinea; *h^C^*, conspicua; *h^Sp^*, spectabilis; *h^A^*, axyridis). The phylogenetic relationships among these alleles were inferred from nucleotide sequences in the conserved region of the first intron of *h*(Ando et al., 2018). *Coccinella septempunctata* (*C. sep*) was used as an outgroup. **(B)** Representative elytra of control (GFP RNAi-treated) females and males, and quantification of black spot size. The area of each black spot (y-axis) was measured and normalized to the total elytral area. In the horizontal axis, “F” and “M” indicate sex (F: female; M: male), and numbers 1 to 10 correspond to black spot positions illustrated in the schematic. Differences in spot size between sexes were compared using the Brunner-Munzel test, and p-values were adjusted by the Holm method. Asterisks indicate statistical significance: *, p < 0.05; **< 0.01; ***< 0.005. Exact p-values are shown when p ≥ 0.05.

In the present study, we investigated the role of *dsx* in the formation of sexual dimorphism in the elytral color pattern of *H. axyridis*. In insects, *dsx* regulates the formation of sexual dimorphism by expressing sex-specific isoforms (*dsxM* in males and *dsxF* in females), which are generated through sex-specific splicing triggered by species-specific sex-determination cues (Verhulst & van de Zande, 2015). DsxM and DsxF proteins have two conserved domains: a common DNA-binding domain (Doublesex and Mab-3 [DM] domain) in the N-terminal region and an oligomerization domain (OD) diverged between sexes in the C-terminal region. These two proteins bind to the same DNA motifs in the genome; however, sex-specific cofactors, which bind to Dsx protein via the distinct sex-specific OD domains, are thought to be recruited to the target genes and activate sex-specific gene regulatory networks associated with sex-specific differentiation (e.g. binding of Intersex protein to DsxF) (Yang, Zhang, Bayrer, & Weiss, 2008).

Here, we investigated the role of *dsx* in the sexual dimorphism formation of the forewing color pattern in the *h* allele (Red-nSpots) of *H. axyridis*. Using RNA interference (RNAi), we found that *dsx* regulates the formation of sexual dimorphism primarily through male-specific modulation of black spot size. Moreover, mRNA-seq analysis suggested that such modulation is mediated primarily through the downregulation of *h*. In addition, the bioinformatics analysis focusing on open chromatin regions and Dsx binding motifs across the four major *h* alleles revealed that the ratio of the open chromatin regions with and without Dsx binding motifs was decreased in the three derived alleles other than the Red-nSpot allele. Based on the results, we discuss how secondary sexual traits have evolved within the species *H. axyridis*.

## Materials and methods

### Ladybug husbandry and strains

Laboratory stocks of the *H. axyridis* were derived from field collections in Japan. Ladybug adults were reared at 25 °C and usually fed on an artificial diet (Niimi, Kuwayama, & Yaginuma, 2005) or on the pea aphid *Acyrthosiphon pisum* for egg collection. The pea aphids were cultured on the broad bean *Vicia faba* seedlings at 20 °C and 60% RH. An experimental strain with one of the Red-nSpots (succinea) class of the *h* allele was isolated by crossing individuals with Red-nSpots alleles and individuals homozygous for the Black-2Spots allele, and then selecting *h* homozygotes from the siblings. Moreover, adults with larger spots were selected for five generations. For experiments, larvae hatched from a clutch of eggs were reared in a plastic cup with a water reservoir and fed aphids until they reached the prepupal stage. Pupae were reared in incubators at 17°C or 25°C until eclosion.

### BLAST search for *doublesex* orthologs

The tblastn program (Camacho et al., 2009) was used to search for the ortholog of *doublesex*. An amino acid sequence of *Tribolium castaneum* Dsx protein (GenBank: NP_001345539.1) was used as a query against an *H. axyridis* cDNA database derived from pupal elytra at 80 h after pupation (80h AP).

To determine the exon-intron structures of the identified genes, we performed a blastn search on the *H. axyridis* genome database using the obtained *H. axyridis dsx* cDNA sequence as a query. The exon-intron structures of the male- and female-specific *dsx* isoforms were determined using sex-specific isoforms found in the cDNA database derived from male and female pupal elytral RNA-seq data.

To design the target region of RNA interference in the common region between *dsxM* and *dsxF*, we performed the NCBI Conserved Domain Search (Wang et al., 2023) and sequence alignment with the *T. castaneum dsx* using the obtained *dsxM homolog* sequences as a query.

### Molecular phylogenetic analysis

To test whether the *dsx* homologs identified in *H. axyridis* were orthologs of *dsx* in *D. melanogaster*, we performed molecular phylogenetic analysis using Dsx orthologs in insects and other DMRT family proteins. We used amino acid sequences of Dsx and Dmrt99B from Zygentoma, Coleoptera, Lepidoptera, and Diptera, as well as those of Dmrt1 from four vertebrate species, as the outgroup (Table S1).

We aligned the amino acid sequences using the MAFFT program (Katoh & Standley, 2013) (version 7) with the -linsi option (to use an accuracy option, L-INS-i) and removed poorly aligned regions using the TrimAl program (Capella-Gutiérrez, Silla-Martínez, & Gabaldón, 2009) (version 1) with the -gappyout option, which uses information based on gaps’ distribution. The alignment figure was generated using Boxshade (https://junli.netlify.app/apps/boxshade/) (version 3.3). The molecular phylogenetic analysis was performed using the aligned sequence as input and the maximum-likelihood method with the IQ-TREE software (Minh et al., 2020). We used the ultrafast bootstrap (UFBoot) and the Shimodaira-Hasegawa approximate likelihood ratio test (SH-aLRT) to evaluate the branch reliability according to the software guideline (Guindon et al., 2010; Minh, Nguyen, & von Haeseler, 2013). We set 1,000 replications in each test. Typically, the branch with the SH-aLRT ≥80% and the UFboot ≥95% is considered reliable (Minh et al., 2020). Therefore, we regarded the branch with both support values exceeding the thresholds as the reliable clade.

### Larval RNAi

The double-stranded RNA (dsRNA) for RNA interference (RNAi) was synthesized based on the T7 RiboMAX Express Large Scale RNA Production System protocol. Briefly, the target region of RNAi was designed within the common exon (the first exon) shared between male- and female-specific isoforms of *dsx* (Fig. 2). For negative control experiments, we used a GFP gene fragment derived from the jellyfish *Aequorea victoria*. The template DNA for dsRNA synthesis was amplified using Polymerase Chain Reaction (PCR) with Q5 DNA polymerase (New England Biolabs, Massachusetts, US) and primers 5′-flanked with T7 promoter sequences (FASMAC, Atsugi, Japan), according to the manufacturer’s instructions (Table S2). The PCR products were separated by 1% agarose gel electrophoresis and purified using NucleoSpin Gel and PCR Clean-up kit (MACHEREY-NAGEL, Düren, Germany). The dsRNA was synthesized using the purified template DNA using the T7 RiboMAX Express Large Scale RNA Production System (Promega, Madison, WI, USA) according to the manufacturer’s instructions with some modifications. The modifications were as follows: in vitro RNA synthesis was performed by incubating at 37°C for 16 hours; annealing was carried out at 65°C for 10 minutes, followed by incubation at room temperature (25°C) for 30 minutes. dsRNA was stored at −80°C until it was used. The microinjection procedure for larval RNAi was performed according to (Niimi et al., 2005). For micro-injections, fine glass needles were prepared by stretching glass capillary tubing (borosilicate glass with filament, O.D.: 1.0mm, I.D.: 0.50 mm, 10cm length, Sutter Instrument, Novato, CA, USA) using a micropipette puller (PC-100, Narishige, Tokyo, Japan) with the STEP1 heating value of 60. Approximately 5.0 µg of the dsRNA was injected into the hemocoel of each final instar larva 1-2 days after molting. The dsRNA was injected at the lateral side of the intersegmental membrane between the second and third thoracic segments using the fine grass needle connected to FemtoJet (Eppendorf, Hunburg, Germany). After injection, 1 to 4 larvae were reared in each container and fed sufficient amounts of aphids at 25 °C until pupation. Prepupae or pupae were incubated at 17°C to minimize the influence of heat-dependent phenotypic plasticity in the Red-nSpots strains (Michie, Mallard, Majerus, & Jiggins, 2010). After eclosion, adult ladybugs were placed in plastic vials with water reservoirs and fed an artificial diet at 25°C for at least two days to ensure color maturation. Finally, we collected the elytra and heads from the ladybugs and stored them at −30°C until phenotypic analysis was performed.

**Fig. 2.**
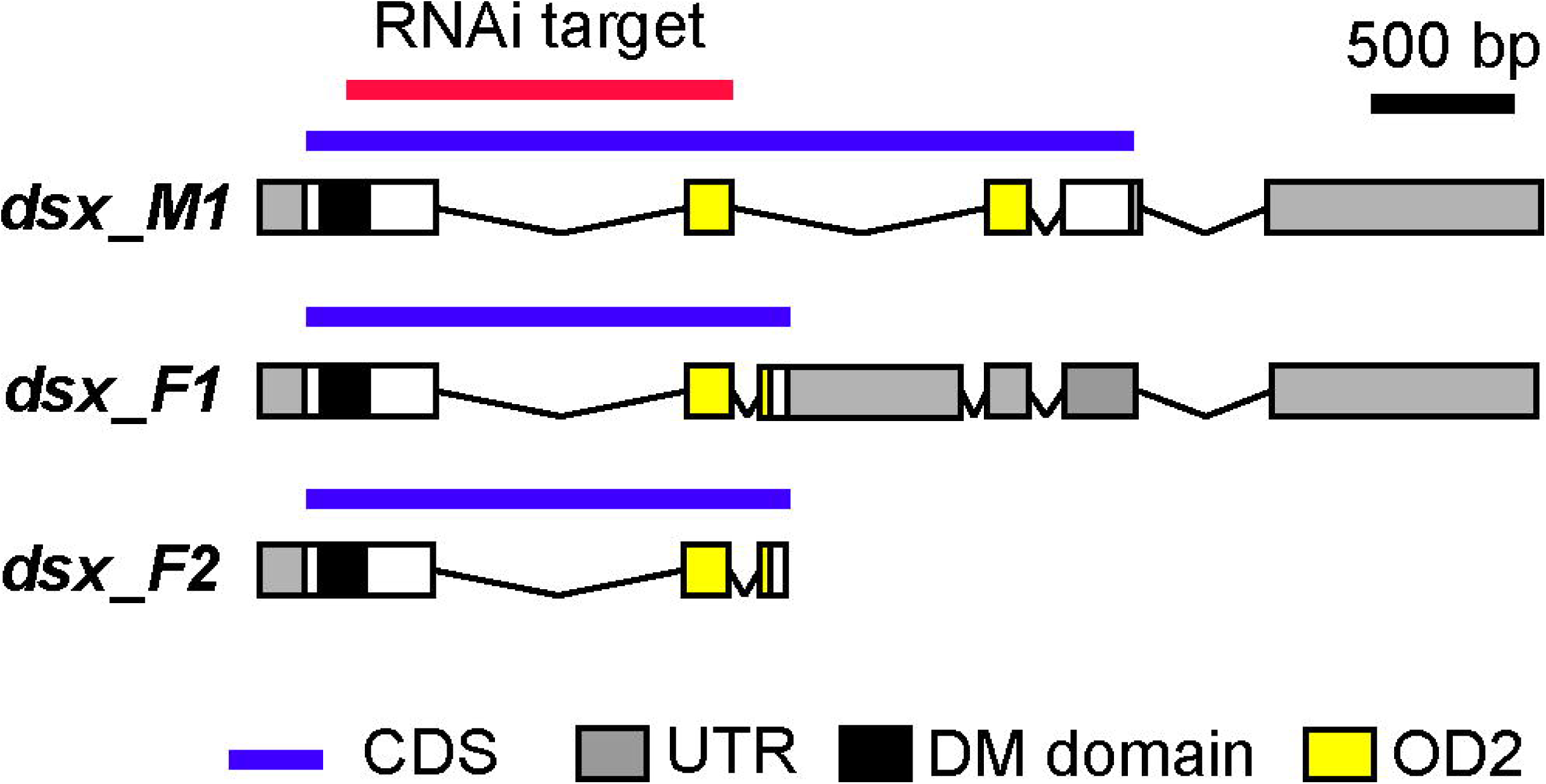
Schematic representation of putative exon-intron structures and sex-specific splicing patterns of *dsx* in *H. axyridis*. The diagram illustrates the predicted exon–intron organization of the *dsx* gene in *H. axyridis*. Red bar: RNAi target region; Purple bar, coding sequence (CDS); Gray box: untranslated region (UTR); Brown box, DM domain; Yellow box, OD2.

### Quantification of black pigmentation in the elytra and head

An isolated elytra or a head was immobilized on a High-Reflectance PTFE sheet (PMR10P1, Thorlabs, Newton, NJ, USA) with a paste eraser for image acquisition. We set up the illumination light with a hand-made dome light apparatus. We collected the elytral or head images using a digital camera (NY-D5500, Nikon, Tokyo, Japan) connected to a stereomicroscope (Stemi 508, Carl Zeiss, Jena, Germany) with an adaptor lens (NTY-1S, Micronet, Kawaguchi, Japan). The shutter speed and ISO were set to 1/10 second and 400, respectively. The contour of the elytra and black spots were automatically extracted using a custom program written in the C language. In this program, the following image processing was performed.

To segment the elytra and spots, we first applied median filtering with a radius of 4 pixels to the resulting RGB images to achieve smoothing. The red channel was used to extract the morphology of the individual spots, while the blue channel was used to extract the overall contour of the forewing. The outline of the entire forewing was obtained by applying Otsu’s thresholding method to the blue channel, followed by segmentation. Similarly, Otsu’s method was also employed to segment the individual spots. Binary images resulting from the segmentation were further processed by contour smoothing using a mathematical morphology operation with a structuring element of radius 8 pixels.

To align elytral contours between individual images, the set of pixels representing the outer shape of each forewing was aligned so that its centroid coincided with the center of the image. Additionally, the images were rotated so that the first and second principal components, obtained through principal component analysis (PCA), were aligned with the X- and Y-axes, respectively.

To automatically localize the averaged characteristic ten-spot patterns in each image, it was necessary to estimate the average position of each spot. For this purpose, the size of the forewing shapes was first normalized. Specifically, the bounding rectangle enclosing each forewing was computed, and the images were scaled so that all specimens had identical bounding rectangle dimensions (width and height). Using the resulting aligned and scaled images, an average image was generated. Based on this image, an arbitrary threshold was determined to allow clear separation of the ten spots, and segmentation was performed. This procedure enabled estimation of the average position and shape of each spot.

We performed constrained watershed segmentation to split fused spots. The appearance of spots varies in size and shape across specimens, and in some cases, multiple spots are fused into a single region. Although watershed transformation is effective for separating such fused spots, it may result in over-segmentation when the distance transform image contains multiple peaks due to complex spot morphologies. To address this issue, the average image of the spots was first segmented and labeled. The peaks in the distance-transformed image were then assigned to one of the labeled spot regions from the average image. This information was used as a constraint during watershed transformation, thereby reducing over-segmentation.

Using the extracted contours of the forewing and its constituent spots, we calculated the areas of the blackspots (Table S3). For the head melanization, the degree of melanization of the frons (the frontal plate of the head) was classified into three levels: 0 = no melanization, 1 = melanized limited to the area below the compound eyes, and 2 = melanization extending above the lower edge of the compound eyes.

### Scoring of abdominal morphological defects in *dsx* RNAi mutants

To score the morphological defects in the sexual dimorphism of the terminal end of the abdominal segment (hypopygium) in *H. axyridis*, we investigated the hypopygial morphology using scanning electron microscopy. The posterior parts of the adult body (half of the thoracic segments and abdominal segments) were dissected with forceps and mounted on a pedestal with its ventral side up using double-sided carbon tape. The samples were analyzed using a tabletop scanning electron microscope, Miniscope (TM4000PlusII, Hitachi High-Tech Corporation, Tokyo, Japan). To assess the morphological defects in the hypopygium, the observer scored the specimen as male or female without knowing its sex or treatment information. After scoring, individuals whose hypopygial morphological score was consistent with the specimen’s sex information were considered normal. The individuals whose morphology was inconsistent were considered abnormal.

### Statistical analysis

All statistical analyses were performed using R (R Core Team, 2023) (version 4.3.2). For continuous data with potentially unequal variances and non-normal distributions (elytral spot area), the Brunner-Munzel test was applied using the brunnermunzel R package (https://CRAN.R-project.org/package=brunnermunzel) (version 2.0). Fisher’s exact test was used to evaluate differences in categorical variables between groups (classification of head pigmentation and hypopygial morphology as normal or abnormal). When the contingency table exceeded 2×2, we estimated the p-values (simulate.p.value = TRUE, B = 10000) using a Monte Carlo simulation with 10,000 replicates. Multiple testing corrections were applied using the Holm method where applicable. A p-value < 0.05 was considered statistically significant.

### Identification of sex based on PCR

The sexes of the *H. axyridis* were identified by genomic PCR using the Y chromosome-specific primers (Gotoh, Nishikawa, Sahara, Yaginuma, & Niimi, 2015) (Table S2 #3, #4). Crude extract samples for PCR were collected from each individual using GenCheck DNA Extraction Reagent (FASMAC, Kawaguchi, Japan) according to the manufacturer’s instructions. PCR was performed using the DNA polymerase, KOD FX Neo (TOYOBO, Osaka, Japan) with the following cycling conditions: 94°C for 2 min, followed by 40 cycles of 98°C for 10 sec, 60°C for 30 sec, 68°C for 8 sec, and 68°C for 2 min. The PCR products were separated by 1% agarose gel electrophoresis.

### mRNA sample preparation and sequencing

To collect elytral total RNA from the RNAi-treated pupae, the final instar larvae were injected with 5 µg of dsRNA targeting *dsx* or *GFP* within two days after molting and reared as described above. Pupation timing was monitored at 25°C under full light conditions using LiveCapture3 software (https://lc3.daddysoffice.com/livecapture3/) (v3.5) and a color CMOS camera (1080P-USB 2.0, Vsightcam, Hangzhou, China).

Pupal elytra were dissected in Dulbecco’s phosphate-buffered saline (D-PBS) at 80 h after pupation (80 h AP). Three biological replicates were prepared for each condition (Male *GFP* RNAi, Female *GFP* RNAi, and Male *dsx* RNAi). The collected samples were snap-frozen in liquid nitrogen and stored at −80°C until use. Total RNA was extracted from each sample (two elytra per sample) using the RNeasy Micro kit (Qiagen, Venlo, the Netherlands) according to the manufacturer’s instructions with the on-column DNase I treatment protocol. The total RNA was quality-checked with 1.0% agarose gel TBE/formamide electrophoresis and quantified using the QuantiFluor RNA system and Quantus Fluorometer (Promega, Madison, WI, USA). The library preparation and the sequencing analysis were performed at the ASHBi SingAC core facility at Kyoto University. Briefly, each sequencing library was prepared using 100 ng of total RNA as input, NEBNext Poly(A) mRNA Magnetic Isolation Module, NEBNext Ultra II Directional RNA Library Prep Kit, NEBNext Multiplex Oligos for Illumina (New England BioLabs, Ipswich, MA, USA), and Biomeck i7 automated workstation (Beckman Coulter, Brea, CA, USA). The 50 bp paired-end sequence data were obtained using the NovaSeq 6000 system (Illumina, San Diego, CA, USA).

### Assay for Transposase-Accessible Chromatin sequencing (ATAC-seq)

Laboratory strains with four different color pattern alleles (Red-nSpots, Black-2Spots, Black-4Spots, and Black-nSpots) were used for ATAC-seq analysis. Pupal forewings at 80 h AP were dissected and homogenized in a 1.5 ml microcentrifuge tube using a plastic pestle (BioMasher-II; Nippi, Tokyo, Japan) with 200 µl of Lysis Buffer (10 mM Tris-HCl, pH 7.4, 10 mM NaCl, 3 mM MgClC, 0.1% Triton X). The nuclear suspension was filtered through a polyester mesh filter (pore size: 74 µm; Costar #3477, 12-well) (Corning, NY, USA) and centrifuged using a swing rotor (800 × g, 4°C, 10 min). After removing the supernatant, the pellet was resuspended in 25 µl of D-PBS. Nuclear concentration was determined by mixing an aliquot (2.5 µl) of each sample and an equal volume of Trypan Blue staining solution and counting the cell nuclei with a hemocytometer under a compound microscope (Axiovert 200M, Carl Zeiss, Oberkochen, Germany). Approximately 0.5–1 × 10^5^ nuclei were used per ATAC-seq library preparation. The nuclei were subjected to transposition with Tn5 Transposase at 37°C for 30 min using Nextera DNA Library Prep Kit (FC-121-1030; Illumina, CA, USA). DNA was purified using the MinElute PCR Purification Kit (Qiagen, Venlo, Netherlands). Library preparation was conducted according to Buenrostro et al. (2015) (Buenrostro, Wu, Chang, & Greenleaf, 2015). Library quality was assessed using the High Sensitivity DNA kit (Agilent Technologies, CA, USA) and the KAPA Library Quantification Kit (Illumina/Universal) (KAPA Biosystems, MA, USA). Sequencing was performed on NextSeq 550 and HiSeq X systems (Illumina, San Diego, CA, USA) (Table S4).

### mRNA-seq data analysis

The quality of the sequencing data was verified using FastQC software (https://www.bioinformatics.babraham.ac.uk/projects/fastqc/). After trimming adapter sequences and low-quality sequences using Cutadapt software (Martin, 2011), the FASTQ sequence data were mapped to the latest *H. axyridis* reference genome (Chen et al., 2021) (ZJU-BJ) using hisat2 software (Kim, Paggi, Park, Bennett, & Salzberg, 2019) with the default parameters. The potential transcription units (the GTF format files) were constructed from each sample and merged with the previously annotated transcription units (Chen et al., 2021) (ZJU-BJ) using the StringTie software (Pertea et al., 2015). The read counts were quantified based on the merged transcription units using the StringTie software.

The count data were normalized between samples for each comparative analysis using the iDEGES method in the TCC package (Sun, Nishiyama, Shimizu, & Kadota, 2013) with three iterations of DESeq2-Trimmed Mean of M-values (TMM) normalization. The differentially expressed genes (DEGs) were identified using the Wald test, followed by fitting the data with a negative binomial generalized linear model (GLM) using the deseq2 method in the TCC package (Sun et al., 2013).

Gene Ontology (GO) terms were assigned to each gene using the eggNOG-mapper2 software (Cantalapiedra, Hernández-Plaza, Letunic, Bork, & Huerta-Cepas, 2021). The GO enrichment analyses were performed using the topGO R package (Alexa & Rahnenfuhrer, 2025) and the DEGs identified in the above analyses as input. We employed the “elim” algorithm and Fisher’s exact test in the topGO package. False discovery rates were calculated using the p.adjust function in the base R package, employing the Benjamini-Hochberg procedure.

The Venn diagram illustrating the inclusion relations of DEGs across different RNAi treatments was created using the Intervene Shiny App (Khan & Mathelier, 2017). The graphs associated with the normalized count data in the mRNA-seq analysis were generated using a custom R script that utilized the following R packages: ggplot2 (https://cran.r-project.org/package=ggplot2), tidyverse (https://cran.r-project.org/package=tidyverse), stringR (https://cran.r-project.org/package=stringr), ggbeeswarm (https://cran.r-project.org/package=ggbeeswarm), and patchwork (https://cran.r-project.org/package=patchwork).

### Motif scanning for the Dsx transcription factor

We searched for the putative Dsx binding motifs in the genomic regions surrounding the gene *h* using the Possum program (https://bu.wenglab.org/possum/). The public genomic scaffold data of the *h* locus from four different alleles of color patterns (Red-nSpots, Black-2Spots, Black-4Spots, and Black-nSpots) in *H. axyridis* were used for the analysis (GenBank accession numbers: Red-nSpots, LC269048.1; Black-2Spots, LC269055.1; Black-4Spots, LC269054.1; Black-nSpots, LC269053.1). The Position Specific Scoring Matrices calculated from ChIP-seq (Chromatin ImmunoPrecipitation-sequencing) analysis in *Drosophila melanogaster* (JASPER: MA1836.1, Fig. 5A) were used as query data. We set the threshold for log-likelihood ratio scores to 5 for detecting sequence fragments similar to the query matrix data of Dsx binding sites.

### ATAC-seq data analysis

The raw Assay for Transposase Accessible Chromatin-sequencing (ATAC-seq) read data used for the analysis were obtained from pupal elytral tissues at 80 h after pupation in four different color pattern types of *H. axyridis*: Red-nSpots, Black-2Spots, Black-4Spots, and Black-nSpots. Three biological replicates for each sample were included in the dataset. We used the latest version of the scaffold-level genome assembly of Black-2Spots *H. axyridis* as the template reference genome (ZJU-BJ (Chen et al., 2021)) for mapping the ATAC-seq data. We swapped the genomic regions surrounding the *h* gene (±500 kb) with the corresponding DNA sequences from different *h* alleles (the same GenBank datasets used in *Motif scanning for the Dsx transcription factor* above).

First, the adaptor sequences were trimmed from the data using the cutadapt software (Martin, 2011) (ver. 4.9). The cropped reads were mapped to the corresponding reference genome sequences using bowtie2 software (Langmead & Salzberg, 2012) (ver. 2.5.3) with --local and -very-sensitive-local options. Duplicate reads were removed using Picard software (https://broadinstitute.github.io/picard/) (ver. 3.3.0) with the - REMOVE_DUPLICATES true option. ATAC-seq peaks were called using macs3 software (Zhang et al., 2008) (ver. 3.0.0b3) with –nomodel, --shift −75, --extsize 150, f - BAMPE, -g 430000000, and -B options. We validated the quality of each ATAC-seq data by calculating the FRiP (Fraction of all mapped Reads that fall into the called Peak regions) score according to the ENCODE project guidelines: ideal > 0.3; acceptable > 0.2 (https://www.encodeproject.org/atac-seq/). All the samples satisfied the criteria of the ENCODE project. We extracted from the obtained peaks those with a q-value <= 0.001. Of the three samples in each allele, two samples with higher FRiP scores were selected to determine the reproducible ATAC-seq peaks. We extracted such peaks using the bedtools intersect program (Quinlan & Hall, 2010) with -wa and -wb options.

## Results

### The function of *dsx* in the sexually dimorphic color pattern formation in *H. axyridis*

As previously reported (Hosino, 1942, 1948; Knapp & Nedvěd, 2013), the *h* allele (the Red-nSpots) used in the present study exhibited significantly larger black spots in females than in males (Fig. 1B). Here, we focused on the molecular basis of this sexual dimorphism. We aimed to test the role of the master regulator of insect sex differentiation, *dsx,* in the formation of sexually dimorphic elytral color patterns of *H. axyridis*. We first identified a *dsx* homolog and male- and female-specific isoforms (Fig. 2, *dsx_M*, and *dsx_F1,2*) in the pupal elytral mRNA-seq data by the BLAST search, using *Tribolium castaneum dsx* ortholog as a query. We also performed conserved domain search (NCBI) and sequence alignment with *T. castaneum* Dsx to infer the DM domain in the first exon, OD2 in the second exon, and the terminal region of the female-specific OD2 in the third exon (Fig. 2). To confirm orthology, we conducted molecular phylogenetic analyses using the coding amino acid sequence of *H. axyridis dsx* homolog and those of DM domain-containing genes (Dsx, Dmrt99B, and Dmrt1) from apterygote, holometabolan, and vertebrate databases (Table S1 and Fig. S1). The molecular phylogenetic tree showed that the *H. axyridis dsx* clustered within the insect Dsx clade and formed a Coleoptera-specific subclade with *T. castaneum* Dsx (Fig. S2), consistent with transcriptome-based insect phylogenies (Misof et al., 2014). These results confirmed that the *dsx* homolog in *H. axyridis* is an ortholog of *Drosophila dsx* and hereafter referred to as *dsx*.

To determine whether *dsx* is expressed in pupal elytra during color pattern prepatterning at 80 hours after pupation (80 h AP) (Ando et al., 2018), we reanalyzed public mRNA-seq datasets from male pupal elytra carrying the Black-2Spots allele (*h^C^*) at 24, 72, and 96 h AP (DDBJ: DRR092246-DRR092257). *dsxM* expression was detectable as early as 24 h AP and peaked around 72 h AP. Notably, *dsx* was expressed in both red and black regions of the elytra (Fig. S3).

To assess the functional role of *dsx* in elytral color patterning, we performed larval RNAi knockdown experiments and measured the size of the black spot in adults. In males, *dsx* RNAi significantly increased spot size compared to the *GFP* RNAi control (Fig. 3A), yielding spot sizes comparable to those of *GFP* RNAi control females (Fig. 3C). In contrast, *dsx* RNAi had no significant effect on spot size in females (Fig. 3B). These findings indicate that the sexual dimorphism in black spot size associated with the *h* allele is primarily due to *dsxM*-mediated suppression of black pigmentation in males.

**Fig. 3.**
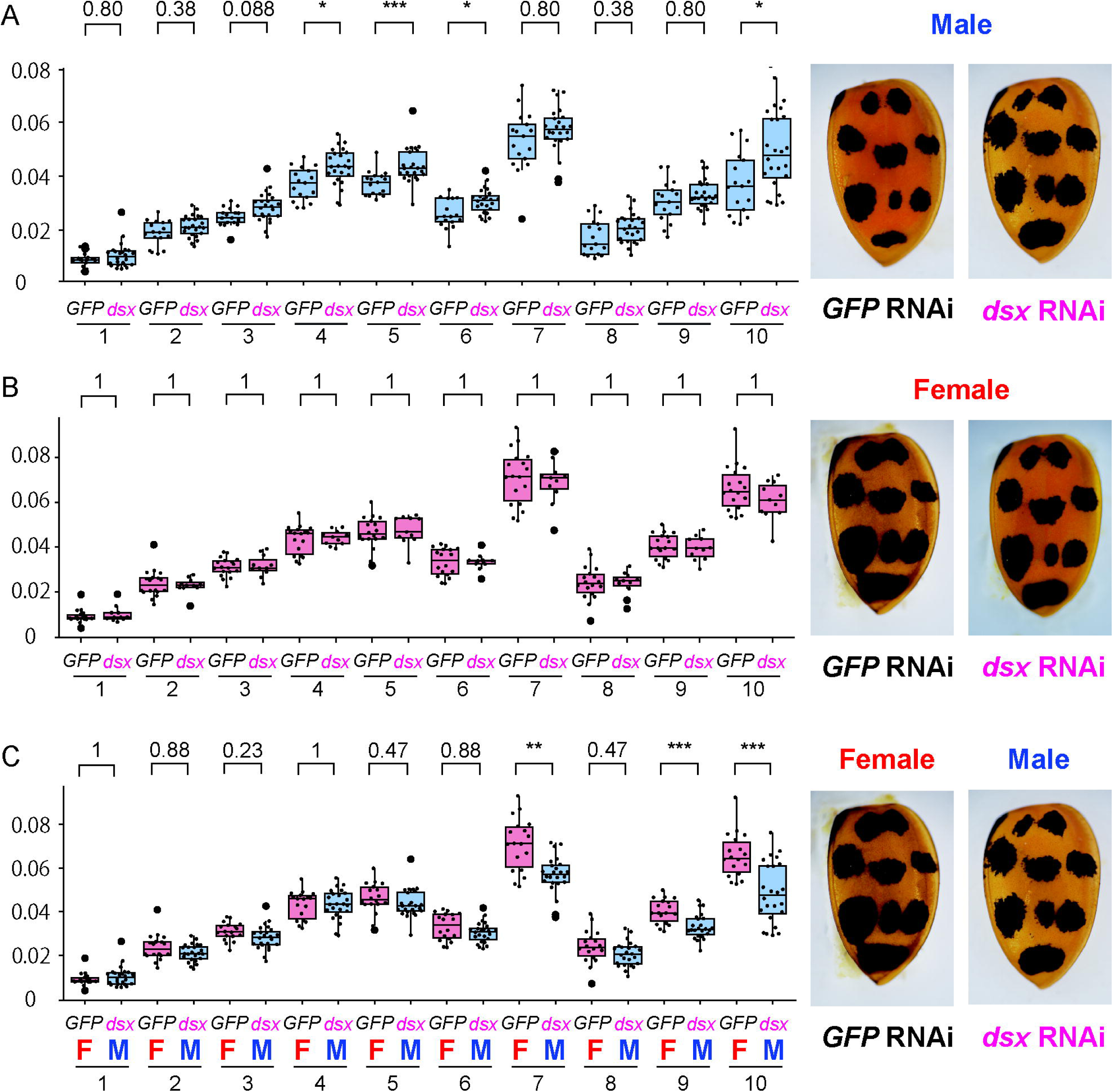
*dsx* knockdown phenotypes in the elytral color pattern. **(A-C)** Comparison of black spot sizes in RNAi-treated individuals: **(A)** between *dsx* RNAi males and control (*GFP*) RNAi males, **(B)** between *dsx* RNAi females and control (*GFP*) RNAi females, and **(C)** between *dsx* RNAi males and control (*GFP*) RNAi females. Black spot areas were quantified and statistically analyzed as described in Fig. 1. Asterisks indicate statistical significance: *, p < 0.05; **, <0.01; ***, <0.005. Representative elytra of individuals from each RNAi treatment are shown on the right side of each graph.

In addition to the elytral color patterning, we observed defects in other external sexually dimorphic traits in *dsx* RNAi-treated individuals with the *h* allele. In head pigmentation, the frons (the frontal edge of the head) is typically white in males and black in females. This pattern was disrupted in both sexes by *dsx* RNAi (Fig. S4, Table S5), in contrast to the male-specific elytral pigmentation phenotype in the elytra. Another affected trait was the morphology of the terminal abdominal segments (hypopygium). In males, the posterior margin of the fifth abdominal segment is anteriorly indented (Fig. S5, arrow), whereas in females, no such indentation is observed, and a small protrusion is formed in the sixth segment (Fig. S5, arrowhead). This sexual dimorphism was lost in *dsx* RNAi individuals, with both sexes exhibiting a shallow indentation and a protrusion of the sixth abdominal segment (Fig. S5, Table S6). Again, *dsx* was found to contribute to sexual differentiation in both sexes, in contrast to its male-biased function in the elytra. In addition to the above morphological defects, we also observed a decrease in the survival rate of *dsx*-RNAi-treated females (Table S7).

### The developmental genes associated with the differentiation of the male and female elytral color pattern at the pupal stage

To further explore the molecular basis of the *dsx*-mediated sexual dimorphism in the elytra, we conducted mRNA-seq analysis on male and female pupal elytra, as well as on male pupal elytra subjected to *dsx* RNAi (*GFP* RNAi in males and females, and *dsx* RNAi in males). To examine the regulatory relationship between *dsx* and the key color patterning gene *h*, we focused on 80 h AP, the developmental stage when *h*-dependent prepattern is established and pigment cell differentiation initiates (Ando et al., 2018).

We first compared transcriptomes of wild-type male and female pupal elytra (male and female *GFP* RNAi) to identify differentially expressed genes (DEGs) between the sexes. Hierarchical clustering of the transcriptome data suggested that gene expression profiles of elytral tissues at 80 h AP were only minimally differentiated between samples (Fig. S6A). Only a small fraction of *H. axyridis* genes was differentially expressed at this stage (94 genes, 0.43% = 94/22,810) (Chen et al., 2021) (Fig. S6B). GO enrichment analysis revealed that DEGs between sexes were significantly enriched for GO terms such as GO:0045433 (male courtship behavior, veined wing-generated song production) and GO:0043009 (chordate embryonic development). Notably, transcription factor genes associated with these GOs included orthologs of *dsx* and the color patterning gene *h*. We further analyzed all transcription factor genes included in the DEG set. Five transcription factors were significantly differentially expressed between female and male elytra at 80 h AP (Fig. 4, Male GFPi; Female GFPi): *dsx*, *h*, *taxi*, and *crooked legs* were downregulated in males, whereas *MADF* was upregulated. These results suggest that the *dsx* may regulate these transcription factors, including *h*, in the pupal elytra. Collectively, our findings implied that a small set of transcription factor genes, including the *h* gene, is differentially expressed between males and females to form sexual dimorphism of elytral color patterns at this prepatterning stage.

**Fig. 4.**
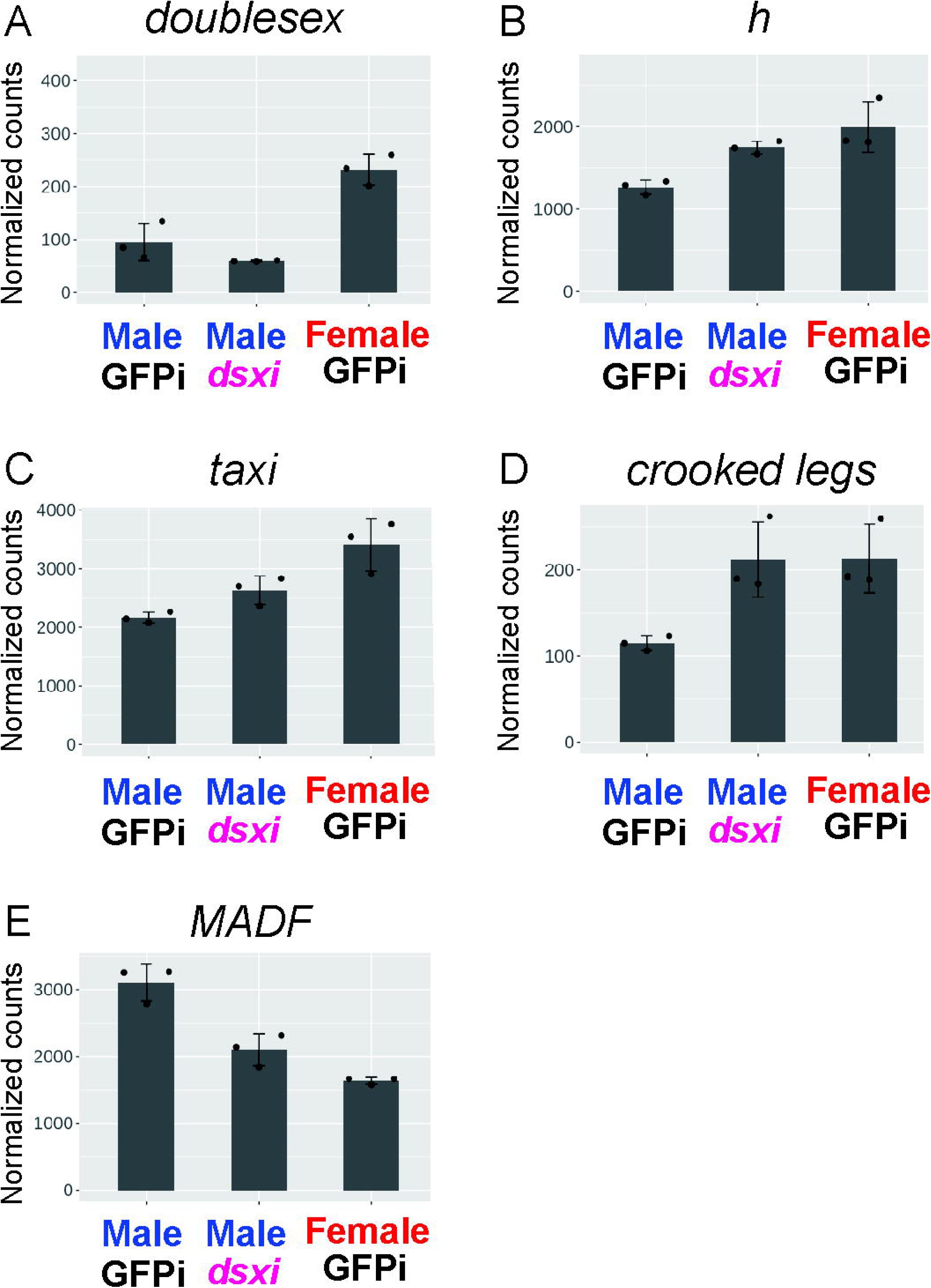
Expression levels of *dsx* and transcription factor genes differentially expressed between sexes. **(A-E)** Normalized expression levels of five transcription factor genes were extracted from transcriptome data and plotted in each graph: **(A)** *dsx*, *h*, **(C)** *taxi*, **(D)** *crooked legs*, and **(E)** *MADF*. Male GFPi: control *GFP* RNAi-treated males. Male GFPi: control *GFP* RNAi-treated females. Male dsxi: *dsx* RNAi-treated males.

### The developmental genes regulated by *dsx* in the sexually dimorphic elytral color pattern formation

To test the above potential regulatory relationship between *dsx* and the four transcription factor genes, we compared the transcriptomes of pupal male elytra treated with *dsx* RNAi and *GFP* RNAi (negative control). Among the four transcription factor genes, *h* and *MADF* showed significantly differential expression in the *dsx* RNA treatment (*h*: upregulation; *MADF*: downregulation) (Fig. 4B, E; Male GFPi, Male dsxi). The expression levels of the other two transcription factors (*taxi* and *crooked legs*) were higher in the *dsx* RNAi treatment, although the differences were not statistically significant (Fig. 4C, D; Male GFPi, Male dsxi). In this experiment, the expression level of *dsx* was also lower in the *dsx* RNAi treatment, although the difference was not statistically significant at this stage (Fig. 4A; Male GFPi, Male dsxi), implying this effect is likely due to the cumulative impact of *dsx* RNAi from the larval to pupal stages. These results suggest that *dsx* regulates at least *h* and *MADF* in the male elytra at 80 h AP.

To further investigate the gene regulatory module downstream of *dsx*, we performed GO enrichment analysis on the differentially expressed genes (DEGs). Among the genes upregulated in male *dsx* RNAi compared to the control (GFP RNAi), significantly enriched GO terms included GO:0030513 (positive regulation of BMP signaling pathway) and GO:0046332 (SMAD binding), suggesting that genes associated with the TGF-β/BMP signaling pathway are negatively regulated by *dsxM* during black spot suppression in male elytra (Table S8). Genes contributing to these GO terms included homologs of *h*, *Daughter against decapentaplegic*, and *magu* (Fig. S7). On the other hand, significantly enriched GO terms among the downregulated DEGs included GO:0007026 (negative regulation of microtubule depolymerization) and GO:0044304 (main axon), implying that *dsxM* positively regulates genes involved in neuronal differentiation in the forewing.

Notably, from these analyses, the key color pattern regulatory gene *h* was identified as a downstream factor of *dsx*. These findings suggest that the regulatory interaction between *dsx* and *h* plays a crucial role in governing sexual dimorphism of the elytral color pattern.

### Open chromatin regions and putative Dsx binding motif in the first intron of the *h* alleles with four different color patterns

To explore the direct regulatory relation between *dsx* and *h*, and its association with the loss of sexual dimorphism in *H. axyridis*, we investigated the distribution of cis-regulatory sequences and Dsx binding motif at the *h* gene in the pupal elytral tissue. Previous genomic analysis revealed that the non-coding regions surrounding the 100 kb-scale first intron of the *h* gene are highly diverged among Red-nSpots (*h*), Black-nSpots (*h^A^*), Black-4Spots (*h^Sp^*), and Black-2Spots (*h^C^*) alleles in *H. axyridis* (Ando et al., 2018; Gautier et al., 2018). In this region, distinct transcription factor-binding motifs are enriched in each allele (Ando et al., 2018).

To estimate the active cis-regulatory elements in the pupal elytra at 80 h AP, we performed ATAC-seq (Assay for Transposase-Accessible Chromatin-sequence), which detects open chromatin regions associated with transcriptional regulatory elements such as enhancers, suppressors, and promoters (Buenrostro, Giresi, Zaba, Chang, & Greenleaf, 2013). In the first intron of the gene *h*, most of the ATAC peaks in each allele were shared between male and female pupal forewings (Fig. S8, 68% in Red-nSpots, 97% in Black-2Spots, 87% in Black-4Spots, and 93% in Black-nSpots). A few sex-specific peaks were detected in all alleles, with slightly more (32%) observed in Red-nSpots. These results imply that the cis-regulatory difference leading to sexually dimorphic expression of the *h* gene in the pupal forewings is dependent mainly on the properties of sex-specific trans regulatory factors (e.g., *dsxM* or *dsxF* isoforms) targeting the same cis-regulatory elements (CREs) shared between the sexes and moderately on fewer sex-specific CREs. We also found that the number of ATAC-peaks in the first intron of the gene *h* increased in lineages with broader black patterns, with counts of 8, 15, 19, and 21 in Red-nSpots (*h*), Black-nSpots (*h^A^*), Black-4Spots (*h^Sp^*), and Black-2Spots (*h^C^*), respectively (Table 1, Fig. 5). This finding suggests that novel cis-regulatory sequences were acquired during the evolutionary process of expanding the expression domain of the *h* gene, which controls the black prepattern of the elytra (Ando et al., 2018; Gautier et al., 2018).

**Fig. 5.**
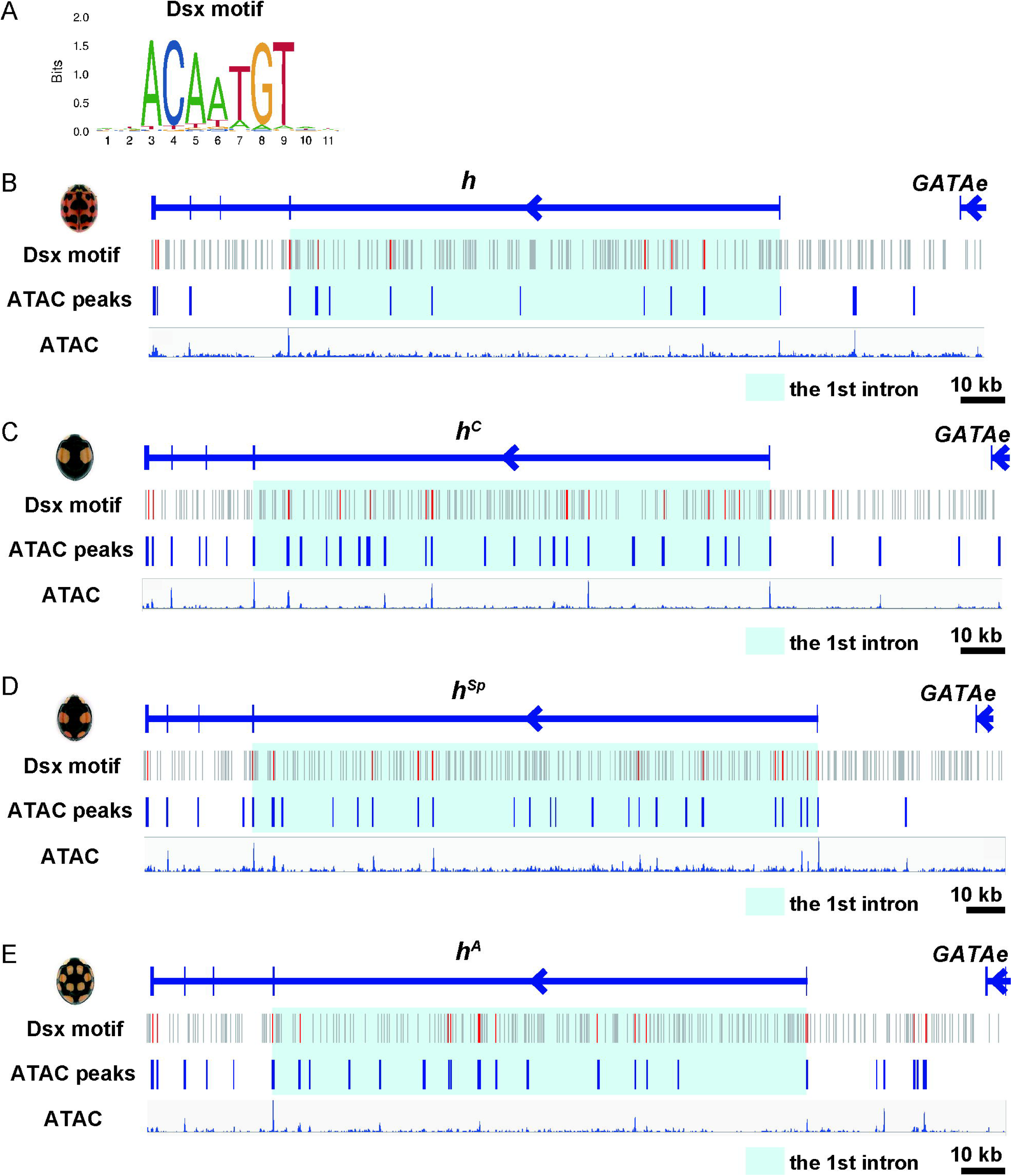
Distribution of Dsx binding motifs and ATAC-seq peaks at the *h* locus in the four major color pattern alleles. **(A)** Sequence logo of the Dsx binding motif in *Drosophila* (JASPAR: MA1836.1). **(B-E)** Each panel depicts the following: **(line 1)** the genomic structure of the *h* allele corresponding to each color morph; **(line 2)** the distribution of Dsx binding motifs (Dsx motif); **(line 3)** the distribution of raw ATAC-seq peaks (ATAC peaks); and **(line 4)** the distribution of raw ATAC-seq reads. **Panels B**-**E** represent the Red-nSpots, Black-2Spots, Black-4Spots, and Black-nSpots alleles, respectively. Grey bars in Dsx motif rows indicate Dsx binding motifs predicted from the *Drosophila* motif sequence; Red bars indicate motifs that overlap with ATAC-seq peaks. Light blue highlights indicate the first intronic region of each *h* allele.

**Table 1.**
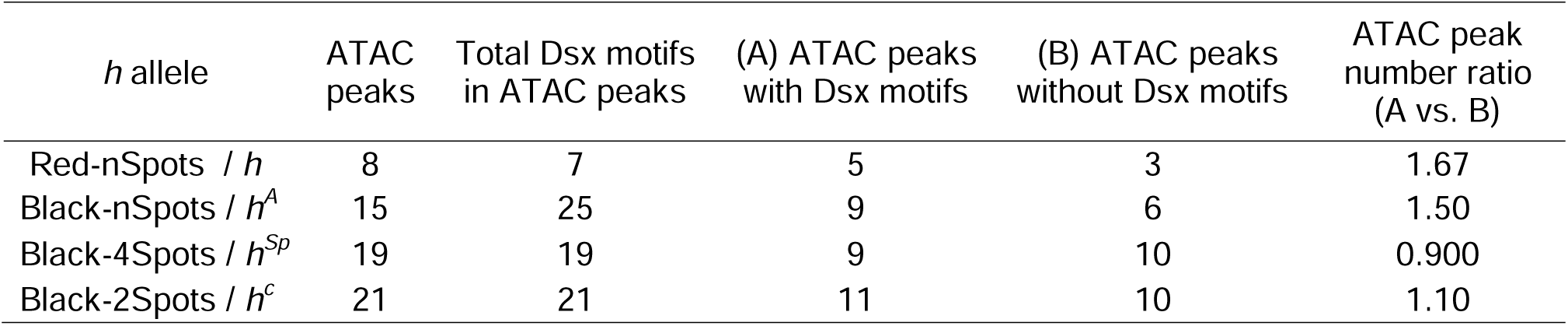
Distribution of ATAC peaks and Dsx binding motifs in the first intron of the four *h* alleles.

To estimate the number of CREs potentially regulated by Dsx, we identified candidate Dsx-binding motifs within accessible chromatin regions, based on a conserved Dsx-binding motif shared between the invertebrate *Drosophila* Dsx and the vertebrate human DMRT1 protein (Murphy, Zarkower, & Bardwell, 2007). We mapped the Dsx binding motif in the open chromatin regions at the *h* locus and compared its distribution between different alleles. The number of open chromatin regions containing Dsx-binding motifs was 5, 9, 9, and 11 in the Red-nSpots, Black-nSpots, Black-4Spots, and Black-2Spots alleles, respectively, with corresponding total motif counts of 7, 25, 19, and 21 (Table 1, Fig. 5). Notably, Red-nSpots, which exhibits clear sexual dimorphism in elytral pigmentation, showed the lowest number of Dsx-binding motifs, suggesting that the absolute number of Dsx-binding motifs is not directly correlated with the presence/absence of elytral sexual dimorphism. In contrast, the ratio of open chromatin regions with versus without Dsx-binding motifs was highest in Red-nSpots (1.67 [5/3]), followed by Black-nSpots (1.50 [9/6]), Black-2Spots (1.10 [11/10]), and Black-4Spots (0.90 [9/10]) (Table 1). These results suggest that a higher proportion of CREs containing Dsx-binding motifs, rather than their absolute number, may be associated with the presence of sexual dimorphism, and that the relative scarcity of such CREs in other alleles may contribute to their loss in pupal elytra at 80 h AP.

## Discussion

In the present study, we investigated the developmental genetic basis of the sexually dimorphic wing color pattern in the harlequin ladybug, *H. axyridis*. We focused on *dsx* as the primary regulatory gene for sexually dimorphic wing color patterning and investigated the pupal elytral development of *dsx* RNAi mutants at the onset of pigmentation. Our RNAi and mRNA-seq experiments revealed that *dsx* mainly inhibits black spot size in males by downregulating the color patterning gene *h*. Moreover, our bioinformatics analysis focusing on the Dsx binding motif and open chromatin regions (enhancer, repressor, and promoter regions) in the pupal elytral development revealed that the Red-nSpots type *h* allele possesses the highest ratio of open chromatin regions with and without Dsx-binding motifs among the four major color pattern alleles. Based on the above findings, we discuss below how *dsx* regulates sexual dimorphism of the elytral color pattern in the Red-nSpots allele and how such regulation was modulated during evolution, along with the loss of sexual dimorphism and the emergence of the novel intraspecific color pattern polymorphism alleles in *H. axyridis*.

### Regulatory mode of sexual dimorphism of pattern regulated by *dsx* in *H. axyridis*: a starting point to understand the evolvability of sexual dimorphisms within a species

We revealed that *dsx* regulates sexual dimorphism of the elytral color pattern in the strain with the Red-nSpots type allele *h*. This regulation, driven by *dsx,* is a typical example of genetic regulation of sexual dimorphism formation in insects. However, elucidating the underlying molecular basis is a crucial first step toward understanding the evolvability of sexual dimorphism and its relationship with the evolution of novel traits within a species. Here, we characterize the regulatory mode of sexual dimorphism formation driven by *dsx* in elytral color pattern formation, as elucidated in the present study.

Regarding the regulation of sexual dimorphism, *dsx* controls elytral pattern dimorphism through a male-specific mode in *H. axyridis*. Generally, in holometabolous insects, *dsx* has contrasting sex-specific functions with one sex-specific isoform (*dsxM* or *dsxF*) inducing sex-specific organ formation and the other (*dsxF* or *dsxM*) suppressing it (e.g., leg sex comb in *Drosophila* (Tanaka et al., 2011), abdominal color pattern in *Drosophila* (Kopp et al., 2000), wing color pattern in swallowtail butterfly (Kunte et al., 2014; Nishikawa et al., 2015), head and thoracic horns in dung beetles and rhinoceros beetles (Ito et al., 2013; Ledón-Rettig, Zattara, & Moczek, 2017; Morita et al., 2019), etc.). Therefore, the male-restricted mode of *dsx*’s regulation in the elytral color formation is a rare example of regulation in holometabolous insects. This difference may stem from the extent to which the sexual dimorphic trait is exaggerated in either males or females. In most cases reported so far, other insects exhibit pronounced sexual dimorphism, characterized by the presence or absence of distinct traits in males and females, respectively (e.g., sex combs, color patterns, horns). In such cases, *dsxM* and *dsxF* must exert strong effects on the transcriptional regulation of target genes to generate sexual dimorphism. In contrast, in the case of sexual dimorphism of elytral color pattern in *H. axyridis*, sexual dimorphism is minimal, and strong effects of both *dsxM* and *dsxF* may not be necessarily required. Since in *H. axyridis*, sexual dimorphism of head pigmentation and abdominal body segment formation is also regulated by the effects of both *dsxM* and *dsxF* (Fig. S5), the above male-restricted regulation does not seem to be due to Dsx protein function specific to *H. axyridis* but is likely due to the characteristics of the regulatory relationship between *dsx* and the target genes in the elytral color pattern formation.

In the following sections, we further discuss the molecular basis of the *dsx*-mediated sexual dimorphism formation in the elytral color patterns and how it influences color pattern evolution.

### The molecular basis of *dsx*-mediated sexual dimorphism formation in the elytral color pattern

We revealed that the male-specific isoform of *dsx* (*dsxM*) primarily functions in the formation of sexual dimorphism in the elytral color pattern and identified the color patterning gene *h* as a crucial downstream target of *dsxM*. Since *h* is a key regulator of pigment cell differentiation, promoting black pigment cell differentiation and suppressing red cell differentiation in *H. axyridis* (Ando et al., 2018), these findings suggest that the regulatory relationship between *dsxM* and *h* is central to the sexual dimorphism formation in the elytral color pattern. The ATAC-seq data and the Dsx-binding motif analysis suggest that the *h* gene in the Red-nSpot allele (*h*) has a high potential to be directly targeted by *dsxM* during pupal elytral development. Additionally, the slight increase in the number of male-specific open chromatin regions in the Red-nSpots allele compared to the other alleles (Fig. S8) implies that these regulatory sequences might also contribute modestly to the sexually dimorphic expression of *h* in the elytra. These male-specific open chromatin regions may be indirectly regulated by *dsxM.* Together, our findings propose the *dsx-* and *h*-mediated molecular platform underlying the formation of sexual dimorphism in the elytral color pattern, which should be further tested through future approaches, such as chromatin immunoprecipitation-based binding assays (Solomon, Larsen, & Varshavsky, 1988) and genome-editing-mediated deletion of candidate regulatory elements (Mazo-Vargas et al., 2022).

In addition to the *h* gene, we identified a small number of regulatory molecules, including transcription factors (*taxi*, *crooked legs,* and *MADF*), and TGF-β-associated genes (*dad* and *magu*), which may act as downstream targets of *dsxM*. These molecules may contribute to the enhancement of sexually dimorphic expression of the *h* gene. Alternatively, they could function downstream of *h*, which itself plays a key role in pigment cell differentiation. Notably, *taxi*, *crooked legs*, *dad,* and *magu* are orthologs of the genes involved in *Drosophila* wing development (D’Avino & Thummel, 2000; Egoz-Matia et al., 2011; Vuilleumier et al., 2010), whereas *h* shows pupal wing expression only in the lineage restricted to ladybugs (Coccinellinae) (Ando et al., 2018; Gautier et al., 2018). These observations suggest that conserved wing development regulators may have been co-opted to enhance the lineage-specific role of *h* in the formation of sexual dimorphism.

In *Drosophila*, not only the prepatterning regulators (e.g., *bric a brac 1* (Williams et al., 2008)) but also effector genes, such as those involved in pigment synthesis (e.g., *tan/CG12120* (Jeong et al., 2008; Luo, Shi, & Baker, 2011)), are direct targets of the sex differentiation cascade in sexual dimorphism formation. However, our mRNA-seq analysis revealed that genes associated with pigmentation were not enriched among the downstream genes of *dsxM* at 80 h AP. This observation is likely due to insufficient expression of pigmentation genes at this developmental stage, making it challenging to determine whether they are regulated by *dsxM*. A similar mode of *dsx*-directed regulation against effector genes may operate in the formation of elytral color pattern dimorphism in *H. axyridis*, which could be uncovered by further analysis focusing on later developmental stages.

### The evolution of sexual dimorphism and Dsx binding motif distribution at the *h* locus

In the present study, we investigated the molecular basis of the sexual dimorphism of the elytral color pattern, which is unique to the basal allele of *H. axyridis*. The findings provide insights into how this regulatory basis may have been modulated during evolution, in parallel with the loss of sexual dimorphism and the emergence of the novel intraspecific color pattern polymorphism in *H. axyridis*. Our analysis pinpointed the significance of changes in *dsx*-dependent regulation targeting *h* during the evolution of sexually dimorphic color patterns.

The pupal elytral cis-regulatory elements at the *h* locus revealed that, among Red-nSpots (*h*), Black-nSpots (*h^A^*), Black-4Spots (*h^Sp^*), and Black-2Spots (*h^C^*) alleles, it was not the absolute number of open chromatin regions with Dsx-binding motifs, but rather the ratio of Dsx motif–containing to motif–free open chromatin regions that was associated with the presence or absence of elytral sexual dimorphism. This observation suggests that the reduction in the ratio of enhancers with and without Dsx-binding site appears to play a crucial role in the evolution of sexual dimorphism in *H. axyridis*. Since the total number of open chromatin regions at the *h* locus was increased in the derived alleles (*h^A^*, *h^Sp^*, and *h^C^*), acquisition of allele-specific CREs might have masked the effect of *dsx*-dependent regulatory elements driving sexually dimorphic *h* expression, even though these CREs are still present in each allele. Such phenotypically masked, cryptic genetic sexual dimorphism—encoded by latent Dsx binding motifs in the cis-regulatory elements—may serve as a hidden reservoir of regulatory potential, enabling the future evolution of novel sexually dimorphic traits. This hypothesis raises an important question to be addressed in future studies aimed at understanding the evolution of sexual dimorphism, as discussed in the previous section, and should be tested through further molecular assays. A recent study on butterfly wing color pattern formation has demonstrated that the synergistic regulation by multiple enhancers drives a precise gene expression pattern in the wing (Mazo-Vargas et al., 2022). This observation suggests that discussion of regulatory evolution should also consider the balance and interaction among multiple cis-regulatory elements.

### Conclusion

In the present study, we investigated the genetic basis underlying the loss of sexual dimorphism in elytral color patterns by focusing on distinct color pattern-type strains of *H. axyridis*. We revealed the crucial male-restricted role of the *dsx* gene in establishing sexual dimorphism in the most basal Red-nSpot allele. From an evolutionary perspective, our findings suggest that the regulatory relationship between *dsxM* and *h* has been modulated during the transition from sexually dimorphic to monomorphic novel elytral color patterns. Our comparative bioinformatics analysis provided a molecular basis of the regulatory model in which the balance between *dsx*-dependent and *dsx*-independent CREs at the *h* locus is essential for understanding the evolution of sexual dimorphism in *H. axyridis*. Overall, this study provides a foundation for understanding the genetic mechanisms underlying the evolution of sexual dimorphism and its relationship to the acquisition of novel color patterns within the species *H. axyridis*.

## Author Contributions

S.Y., T.N., N.H., T.D., and T.A. conceived the project. T.N. selected the *H. axyridis* Red-nSpots strain. S.Y. performed all experiments in this study and analyzed the ATAC-seq data and Dsx motifs. T.A. analyzed the mRNA-seq data. K.K. conducted image analysis and developed a custom program to extract morphological information of elytral black spots. S.M. and T.A. optimized the nuclear extraction protocol for pupal tissues and prepared ATAC-seq libraries. T.A. and K.S. sequenced the ATAC-seq libraries. S.Y. and T.A. wrote the initial manuscripts. All authors reviewed and approved the final manuscript.

## Acknowledgments

We thank Dr. Yuji Matsuoka for his critical comments on the manuscript; Ms. Haruka Kawaguchi for selecting *H. axyridis* strains; Mr. Takeshi Mizuatni for technical support in molecular biology experiments; Mr. Yu Takatani, Ms. Yukiko Sado, Mr. Hiiragi Nishizawa, Mr. Tomoyuki Yokogawa, and Ms. Yurina Takahashi for maintenance of *H. axyridis*; Ms. Yuko Kichise, Dr. Shuji Shigenobu, and Ms. Syoko Ohi for sequencing support for ATAC-seq; Single-cell Genome Information Analysis Core (SignAC) at WPI-ASHBi, Kyoto University, for sequencing support for mRNA-seq; Model Organisms Facility at National Institute for Basic Biology for technical support; and all members of T.D. laboratory and N.H. laboratory for helpful discussion. Computations were partially performed using the supercomputers at the ROIS National Institute of Genetics and the Research Center for Computational Science, Okazaki, Japan (Projects: 24-IMS-C243 and 25-IMS-C328). This research was partially supported by the Cooperative Research Grant of the Genome Research for BioResource, NODAI Genome Research Center, Tokyo University of Agriculture, and the NIBB Collaborative research projects for integrative genomics (Projects: 22NIBB342, 22NIBB448, 23NIBB457, 23NIBB527, and 24NIBB461).

## Data availability

Raw sequencing data were deposited on NCBI (accession numbers: SRR34549913-SRR34549924, SRR34550360-SRR34550368). Raw elytral images used in the morphological analysis were deposited on Figshare (doi: 10.6084/m9.figshare.29653493).

