## Supplementary Information for "Sex-specific *doublesex* regulation targeting the color-patterning gene *h* underlies the evolution of wing sexual dimorphism in the harlequin ladybug *Harmonia axyridis*"

**Supplementary Figure Legends**

**Fig. S1. Multiple alignment of the protein sequences of DMRT-type transcription factors in Insecta and Vertebrata.**

Multiple sequence alignments were generated for the amino acid sequences of Dsx and the other Dmrt family transcription factors. Poorly aligned regions were removed using the TrimAl program. Black boxes indicate conserved residues shared by more than 80% of taxa. The region marked with a red bar corresponds to the DM domain, which is highly conserved among DMRT-type transcription factors.

**Fig. S2. Molecular phylogenetic tree of DMRT-type transcription factors in Insecta and Vertebrata.**

The maximum likelihood (ML) phylogenetic tree was constructed based on the protein sequence alignment shown in Fig. S1. The best hit substitution model was JTT+R2. The proportion of invariable sites is 0.172, and the gamma shape parameter (alpha) was 1.036. Numerical values on each node indicate SH-aLRT and UFBoot support values. Dsx in *H. axyridis* formed a Coleoptera Dsx clade with *T.castaneum* Dsx within the broader Insecta Dsx clade. This result supports that the *dsx* homolog in *H. axyridis* is orthologous to *Drosophila* *dsx*.

**Fig. S3. Developmental expression profile of *dsx* in the pupal elytra.**

*dsx* expression in the red (Red) and black (Black) regions of the pupal wing at different developmental stages (1, 3, and 4 days after pupation [days AP]). *dsx* expression was detected as early as 1 day AP and was observed in both red and black regions of the elytra throughout development.

**Fig. S4.** **Effects of *dsx* knockdown on head pigmentation**

Comparison of the head color in RNAi-treated individuals between *dsx* RNAi and *GFP* RNAi treatments in both sexes. In wild-type individuals, the frons (the frontal surface of the head) is typically white in males and black in females. We found that *dsx* RNAi-treated males exhibited ectopic frons pigmentation, which was absent in wild-type males, whereas *dsx* RNAi-treated females showed reduced pigmentation compared to wild-type females.

**Fig. S5. Effects of *dsx* knockdown on hypopygial morphology**

Comparison of the hypopygial morphology in RNAi-treated individuals between *dsx* RNAi and *GFP* RNAi treatments in both sexes. In wild-type individuals, the posterior margin of the fifth abdominal segment is concave in males (arrow), whereas it is flat in females, with a median protrusion on the sixth segment present only in females (arrowhead). We observed morphological abnormalities in both male and female *dsx* RNAi individuals. Specifically, both *dsx* RNAi males and females showed reduced concavity of the fifth segment (arrow), and a small protrusion on the sixth segment (arrowhead).

**Fig. S6. Comparison of transcriptomes between *dsx* RNAi-treated males and controls (males/females)**

**(A)** Hierarchical clustering analysis of the transcriptome data showing overall similarity or divergence in gene expression among the three groups. **(B)** Venn diagram showing the number of differentially expressed genes (DEGs) between *dsx* RNAi-treated males (dsxi) and control RNAi-treated males (GFPi), and between control RNAi-treated males and females (control RNAi, GFPi).

**Fig. S7. Upregulation of TGF-β-associated genes in *dsx* RNAi in males.**

Normalized expression levels of three genes—**(A)** *h*, **(B)** *dad*, and **(C)** *magu*—were extracted from transcriptome data. Expression profiles are shown for *dsx* RNAi-treated males (dsxi), control RNAi-treated males (Male GFPi), and control RNAi-treated females (Female GFPi).

**Fig. S8. Distribution of Dsx binding motifs and ATAC-seq peaks at the *h* locus in females and males of the four major color pattern alleles.**

**(A–D)** Distribution of chromatin accessibility and Dsx binding motifs at the *h* locus for four representative *h* alleles: (A) Red-nSpots, (B) Black-2Spots, (C) Black-4Spots, and (D) Black-nSpots. Each panel consists of: (line 1) Schematic of the genomic structure. (lines 2-4) ATAC-seq read coverage: (2) female replicate 1, (3) female replicate 2, (4) male. (lines 5–7) Distribution of ATAC-seq peaks: (5) merged peaks from females and males (“All”), (6) peaks shared by both sexes (“common to both sexes”), and (7) peaks that contain Dsx binding motifs (“with Dsx motif”).
