## Supplementary Figure 1 for "Sex-specific *doublesex* regulation targeting the color-patterning gene *h* underlies the evolution of wing sexual dimorphism in the harlequin ladybug *Harmonia axyridis*"

```
                    1 .........10........20........30........40........50........60
Haxy_Dsx            1 --MS--------------DSQEFEIKTDT-------------------------------
Dmel_Dsx            1 M-VS--EE-NWN-SDTMSDSDMIDSKNDVCGG----------------------------
Lcup_Dsx            1 M-VS--EDTNWNSSDTMSDTDMHDSKNDICGG----------------------------
Tcas_Dsx            1 --MS-------------SDSQDFDSKMDV-------------------------------
Bmor_Dsx            1 M-VS--MG-SW----KRRVPDDCEERSEP-------------------------------
Ekue_Dsx            1 M-VS--VG-AW----RRRAPDDCEERSEP-------------------------------
Tdom_Dsx            1 M-MADNQDTNLSSPETVVGVDNSPPEGLENGGL---------------------------
Dmel_Dmrt99B        1 --MS--LP-------SGVDMQNLMSQHPVLGALPP----------AFFLR----------
Lcup_Dmrt99B        1 --MS--LP-------SGVDMQNLMSQHPVLGALPP----------AFFLR----------
Tcas_Dmrt99B        1 --MS--LP------SSGVDMSSLMSQHPVLGAIPP----------AFFLR----------
Bmor_Dmrt99B        1 MEMS--LP-------NGVDVSQLV-QHPVLAALPP-----------FFLR----------
Bmut_Dmrt1          1 --MP--ND-------DAYSKPSAPSEAPQTPGAPPQGKAGG-----------------GG
Drer_Dmrt1          1 --MS--EE-------E--------------------------------------------
Mmus_Dmrt1          1 --MP--ND-------DTFGKPSTPTEVPHAPGAPPQGKAGGYSKAAGAMAGAAGGSGAGG
Xlae_Dmrt1          1 --MQ-NNE-------EPYSKTRNSGQHPS-------------------------------

                                                DM domain (Not trimmed)
                   61 .........70........80........90.......100.......110.......120
Haxy_Dsx           14 -----------NASSTSTNPRTPPNCARCRNHRMKIPLKGHKRYCKYRNCACEKCRLTSE
Dmel_Dsx           28 ---------ASSSSGSSISPRTPPNCARCRNHGLKITLKGHKRYCKFRYCTCEKCRLTAD
Lcup_Dsx           30 ---------ASSSSGSSGTPRTKPNCARCHNHGFKIKLKGHKRYCKYRNCNCEKCRLTAD
Tcas_Dsx           15 -----------NASSTSASPRTPPNCARCRNHRLKIALKGHKRYCKYRTCKCEKCRLTTE
Bmor_Dsx           22 ------------GASSSGVPRAPPNCARCRNHRLKIELKGHKRYCKYQHCTCEKCRLTAD
Ekue_Dsx           22 ------------GATSSGVPRAPPNCARCRNHRLKIELKGHKRYCKYRNCMCEKCRLTAD
Tdom_Dsx           33 ------------GTSSGQNARTPPKCARCRNHRLKIPLKGHKRYCKYRFCNCDKCLLTAE
Dmel_Dmrt99B       30 -------------AAS-ERYQRTPKCARCRNHGVVSALKGHKRYCRWRDCVCAKCTLIAE
Lcup_Dmrt99B       30 -------------AAS-ERYQRTPKCARCRNHGVVSALKGHKRYCRWRDCLCAKCTLIAE
Tcas_Dmrt99B       31 --------------AS-ERYQRTPKCARCRNHGVVSALKGHKRYCRWRDCNCAKCTLIAE
Bmor_Dmrt99B       30 --------------AS-ERYQRTPKCARCRNHGVVSALKGHKRYCRWRDCVCAKCTLIAE
Bmut_Dmrt1         33 GGGGSDSGASGTGAVSGKKSPRLPKCARCRNHGYASPLKGHKRFCMWRDCQCKKCNLIAE
Drer_Dmrt1          6 ---------QTNGSLSIRKPSRMPKCSRCRNHGFVSPLKGHKRFCNWRDCQCQKCRLIAE
Mmus_Dmrt1         50 SGGASGSGPSGLGSGS-KKSPRLPKCARCRNHGYASPLKGHKRFCMWRDCQCKKCSLIAE
Xlae_Dmrt1         20 ------------GVHT-KKSPRLPKCARCRNHGYASPLKGHKRYCMWRDCQCKKCSLIAE

                  121 .........130.......140.......150.......160.......170.......180
Haxy_Dsx           63 RQRVMAMQTALRRAQAQDEAMAKSG-----------------------------------
Dmel_Dsx           79 RQRVMALQTALRRAQAQDEQRALHMHEVPPANPAATTLLSHHHHVAAPAHVHAHHVHAHH
Lcup_Dsx           81 RQRVMALQTALRRAQQQDEQRILQMHEVPPVVHPPPALLKAHYH-------HHHHLQHHL
Tcas_Dsx           64 RQRVMAMQTALRRAQAQDEAMLRSG-----------------------------------
Bmor_Dsx           70 RQRVMAKQTAIRRAQAQDEARARAL-----------------------------------
Ekue_Dsx           70 RQRVMALQTALRRAQAQDEARARS------------------------------------
Tdom_Dsx           81 RQRVMALQTALRRAQAQDEARAAGQ--IP-------------------------------
Dmel_Dmrt99B       76 RQRVMAAQVALRRQQAQEENEAREL-----------------------------------
Lcup_Dmrt99B       76 RQRVMAAQVALRRQQAQEENEAREL-----------------------------------
Tcas_Dmrt99B       76 RQRVMAAQVALRRQQAQEENEAREL-----------------------------------
Bmor_Dmrt99B       75 RQRVMAAQVALRRQQAQEENEAREL-----------------------------------
Bmut_Dmrt1         93 RQRVMAAQVALRRQQAQEE----EL-----------------------------------
Drer_Dmrt1         57 RQRVMAAQVALRRQQAQEE----EM-----------------------------------
Mmus_Dmrt1        109 RQRVMAAQVALRRQQAQEE----EL-----------------------------------
Xlae_Dmrt1         67 RQRVMAAQVALRRQQAQEE----EL-----------------------------------

                  181 .........190.......200.......210.......220.......230.......240
Haxy_Dsx           88 ------------------------AID---------------------------------
Dmel_Dsx          139 AHGGHHSHHGHVLHHQQAAAAAAAAPSAPASHLGGSSTAASSIHGHAHAHHVHMAAAAAA
Lcup_Dsx          134 AEQLHHHHHPHLVDATTSAAVVGAAVP---------------------------------
Tcas_Dsx           89 -----------------------SAVD---------------------------------
Bmor_Dsx           95 ---------------ELGIQPPGLELD---------------------------------
Ekue_Dsx           94 -----------------GVQANGVELD---------------------------------
Tdom_Dsx          108 -----------------GAQS---------------------------------------
Dmel_Dmrt99B      101 -----------------GLLYTSVP-----------------------------------
Lcup_Dmrt99B      101 -----------------GLLYTSVPPR---------------------------------
Tcas_Dmrt99B      101 -----------------GILF---------------------------------------
Bmor_Dmrt99B      100 -----------------NLLYAGQPSS---------------------------------
Bmut_Dmrt1        114 -----------------GISH---------------------------------------
Drer_Dmrt1         78 -----------------GICS---------------------------------------
Mmus_Dmrt1        130 -----------------GISH---------------------------------------
Xlae_Dmrt1         88 -----------------GISH---------------------------------------
```

```
              241 .......250.......260.......270.......280.......290.......300
Haxy_Dsx       91 ----------------------------PHILQ------NTPSPI---------------
Dmel_Dsx      199 SVAQHQHQSHPHSHHHHHQNHHQHPHQQPATQTAL----RSPPHSDHGGSVGPATSSSGG
Lcup_Dsx      161 ----------PHHHHHHHVTH----AAAAAAISTI----RSPPHSDH--SVNGGSSAGGG
Tcas_Dsx       93 ----------------------------PAIMQVP---LKSPSPIH--------------
Bmor_Dsx      107 -------------------------RPVPPVVKAP----RSPMIPP--------------
Ekue_Dsx      104 -------------------------RPDPPAVKTQ----RSPVPP---------------
Tdom_Dsx      112 ----------------------------PVDMESSAGASSIPGPS---------------
Dmel_Dmrt99B  109 -------------------------------GQQN---GSDSATPTP-------------
Lcup_Dmrt99B  111 ---------------------------VPSQNTNE---NTTTATTSQ-------------
Tcas_Dmrt99B  105 ----------------------------PTPAGVV---ADTPGVTAP-------------
Bmor_Dmrt99B  110 ---------------------------AP-------------------------------
Bmut_Dmrt1    118 ----------------------------PIPL---------PSTAEL-------------
Drer_Dmrt1     82 ----------------------------PINL----------SGSDT-------------
Mmus_Dmrt1    134 ----------------------------PIPL---------PSAAEL-------------
Xlae_Dmrt1     92 ----------------------------PIHL---------PIAAEL-------------

              301 .......310.......320.......330.......340.......350.......360
Haxy_Dsx      102 --------------------------------------LLKRKL-------DCDSSSSSQ
Dmel_Dsx      255 GAPSSSNAAAATSSNGSSGGGGGGGGGSSGGG-------------AGGGRSSGTSVITSA
Lcup_Dsx      201 G------------------GNNGGGGGSAGGGVVSSVGAIERNAAALNGMASSSSIASSS
Tcas_Dsx      108 --------------------------------------AIERSL-------DCDSSASSQ
Bmor_Dsx      124 --------------------------------------SAPRSL----GSASCDSVPGSP
Ekue_Dsx      120 ----------------------------------------PRSL----GSASCDSVPGSP
Tdom_Dsx      129 --------------------------------------SLSRE--TVVSAGGCDSSSTSP
Dmel_Dmrt99B  122 --------------------------------------HSPNHSGSGGSGSGGSGQNGVF
Lcup_Dmrt99B  128 --------------------------------------LSPQQTSA----------NGVF
Tcas_Dmrt99B  121 --------------------------------------AIPQ---NSDVGISQLMQRNTF
Bmor_Dmrt99B  112 ------------------------------------------------------------
Bmut_Dmrt1    128 --------------------------------------MVKR---ENSSGNPCLMIESSS
Drer_Dmrt1     91 --------------------------------------LVKN---EAVGEN-VFTLSSGP
Mmus_Dmrt1    144 --------------------------------------LVKR---ENNASNPCLMAENSS
Xlae_Dmrt1    102 --------------------------------------LIKK---EHGGSSSCLMLENSS

              361 .......370.......380.......390.......400.......410.......420
Haxy_Dsx      117 CSPP--------------------------------------------------------
Dmel_Dsx      302 ------------------DHHM--------------------------------------
Lcup_Dsx      244 TAGPPHHPS--------PDHHQQHNNQ---HH-----HHHHP------------------
Tcas_Dsx      123 CSNP--------------------------------------------------------
Bmor_Dsx      142 GVSP--------------------------------------------------------
Ekue_Dsx      136 AVSP--------------------------------------------------------
Tdom_Dsx      149 SSSN------GV------------------------------------------------
Dmel_Dmrt99B  144 HGGIPSPPSDGFEASSTPNHHQQSQQQQQAHHQQHLSHPHQQSQRFNGNESDTDGRTRSE
Lcup_Dmrt99B  140 HAGIPSPPSDGLDSTNNSTHH----------------HTRMPFSNGNSGDSDSEMRIRAE
Tcas_Dmrt99B  140 TASDSSEP----------------------------------------------------
Bmor_Dmrt99B  112 -------------------HH----------------YADMP------------------
Bmut_Dmrt1    147 SSQPPPAS----------------------------------------------------
Drer_Dmrt1    109 PSPASSSA----------------------------------------------------
Mmus_Dmrt1    163 SAQPP-------------------------------------------------------
Xlae_Dmrt1    121 TQTTSTPT----------------------------------------------------

              421 .......430.......440.......450.......460.......470.......480
Haxy_Dsx      121 ---------------------------------------------------PK-------
Dmel_Dsx      306 -----------------TTV-------------------------------PT-------
Lcup_Dsx      270 -------------HPHSSAV-------------------------------PP-------
Tcas_Dsx      127 ---------------------------------------------------PP-------
Bmor_Dsx      146 -------------YAPPPSV-------------------------------PP-------
Ekue_Dsx      140 -------------YAPQPPLS-----------------AP-----------PP-------
Tdom_Dsx      155 ------------VVAVPSRL-------------------------------PP-------
Dmel_Dmrt99B  204 HLSVGFSPTRTELDESPVSKRGAALSNETDQDTGSESGSP--------SSPRPKVAGLFN
Lcup_Dmrt99B  184 RLSMGFSPDRTEV-ESPGSKR-ARLSNETDQDTGSES-AP--------SSPHPKGSNYHN
Tcas_Dmrt99B  148 --------------SSPTSKR-PRINVEDCSLEGSDS-EPEDLKKSRQSSPVP-------
Bmor_Dmrt99B  119 -------------DGSPVQKR-ARITEVC-----STSPSP--------INEQPRDP---K
Bmut_Dmrt1    155 -------------TPSTAAPGPGYSCFFF-------------------------------
Drer_Dmrt1    117 -------------TASPTNLGSRSMLSLS-------------------------------
Mmus_Dmrt1        ------------------------------------------------------------
Xlae_Dmrt1    129 -------------SGSTASSEGKVLIQEI-------------------------------
```

```
                481 .......490.......500.......510.......520.......530.......540
Haxy_Dsx        123 IMRTMSPL-AEPPTTSSSMGAVAQSTD---------------------------------
Dmel_Dsx        311 -PAQSLEGSCDSSSPS---------------PSSTSGAAILPISVSVNR---KNGANVPL
Lcup_Dsx        279 -TAQSVDSSCDSSSPS---------------PSSTSGAISLPVNRKIVPEHHQNGADMSI
Tcas_Dsx        129 AIRKMTPVPAVPSSTSVNIGTIAQSTD---------------------------------
Bmor_Dsx        155 -PPTMPPL-IPPPQPH---------------YWWPGAFPVSPGHVSEQRLS-QEGNIKAV
Ekue_Dsx        152 -PPNMPPL-LPPPQP-------------------------------------------AV
Tdom_Dsx        165 -TARPVKLGIPPSSGH----------------------PAMPPNVTVGE-----------
Dmel_Dmrt99B    256 LTASLSPARTGPPSSP--ESDLDVDSAPDEA---------TPENLSLKKEDSQSPPNTPA
Lcup_Dmrt99B    233 -NGNLSPARTGPPSSP--ESDLDVDSAPDEA---------TPENLSLKKEDSTSPPHTPS
Tcas_Dmrt99B    185 APAPSAPTPSPEPQTSP-DPDLDVEEDTQSE---------APENLSLKKPSSPETPPQPT
Bmor_Dmrt99B    149 TPAPTPPLATPPPRSPS-DHELNVEEELEES-----EPSLTPENLSMKESR-EEMPVKKE
Bmut_Dmrt1      171 -----------PAVTS--RGHVENTPDLVSD---------STYYSSFYQP-----SLFPY
Drer_Dmrt1      133 -----------PAMSS--RGHTDCTSDLMVD---------ASYY-NLYQPT----PYSSY
Mmus_Dmrt1          ------------------------------------------------------------
Xlae_Dmrt1      145 -----------PSITS--RGHMESTSDLVMD---------SPYYSNFYQP-----PLYPY

                541 .......550.......560.......570.......580.......590.......600
Haxy_Dsx        149 -----LLEDCQK-------LLERFKY----------------------------------
Dmel_Dsx        352 -GQDVFLDYCQK-------LLEKFRY----------------------------------
Lcup_Dsx        323 ---DLILDYCQK-------LIEKFGY----------------------------------
Tcas_Dsx        156 -----LLEDCQK-------LLERFKY----------------------------------
Bmor_Dsx        197 -PSETLVENCHR-------LLEKFHY----------------------------------
Ekue_Dsx        167 -SLENLVENCQK-------LLEKFHY----------------------------------
Tdom_Dsx        191 -NVEVLKDSLHA-------LLDMFRL----------------------------------
Dmel_Dmrt99B    305 -ENLHLLRSFSSSHAQGFLPYHHTQFLAAAGLPAHHHP--AAHSPHHQQQQQQQQQQQNH
Lcup_Dmrt99B    281 NDSMHMLRSFNNGNAQGFMPYHHTQFLAAAAASAHSHAQHAQHSPHHPQQPQMP------
Tcas_Dmrt99B    235 ---------------QNFIPYQQFAF----------------------------------
Bmor_Dmrt99B    202 ---------------EEKWENNEFKY---------------------------KMFRRKD
Bmut_Dmrt1      204 -------------------YNNLYNY----------------------------------
Drer_Dmrt1      166 -------------------YSNLYNY----------------------------------
Mmus_Dmrt1          ------------------------------------------------------------
Xlae_Dmrt1      178 -------------------YNNLYNY----------------------------------

                601 .......610.......620.......630.......640.......650.......660
Haxy_Dsx        163 -PWE------------------MMPL------------MHAILKGARADLEEASRRIDEG
Dmel_Dsx        370 -PWE------------------LMPL------------MYVILKDADANIEEASRRIEEA
Lcup_Dsx        339 -PWE------------------MMPL------------MYVILKDAGVDIDEASKRIEEG
Tcas_Dsx        170 -PWE------------------MMPL------------MYAILKDARADLEEASRRIDEG
Bmor_Dsx        215 -SWE------------------MMPL------------VLVIMNYARSDLDEASRKIYEG
Ekue_Dsx        185 -SWE------------------MMPL------------VLVILNYAGSDLDEASRKIDEA
Tdom_Dsx        209 -PLE------------------TLPL------------IYVVLKDARSDVKEASNRIMEA
Dmel_Dmrt99B    362 LPQHHQQQQQQQQQQQRSPIDVLMRV----FPNRRRSDVEQLMQRFRGDVLQAMECMLAG
Lcup_Dmrt99B    335 -PQH-----PSASQQQRSPVDVLMRV----FPNRRRSDVEQLLQRYRGDVLQTMEAMISG
Tcas_Dmrt99B    246 -PPF-----QPQYPAQRSPIDVLMRV----FPGKRRSDVEALLQRCKGDVVQAMEMMVSG
Bmor_Dmrt99B    220 LPEP----AHEESQYQKSPVDVLLKV----FPRRSRQEIEAILARCKGDVVAAMDSMVNG
Bmut_Dmrt1      211 -PQY-----PMALAADSSSGDVGNPLGGSPVKNSLRSLPAPYVPGQTGNQWQMKNS----
Drer_Dmrt1      173 -QQY-----QM--------------------------------PSGNG------------
Mmus_Dmrt1          ------------------------------------------------------------
Xlae_Dmrt1      185 -PPY-----QMAMAAESTSGNDMGGISGPPLKNNHRNHPAAYVPSQSGNQWQMKNR----

                661 .......670.......680.......690.......700.......710.......720
Haxy_Dsx        192 KEA--EI----LLEFCQRIRDK---FQISWRMVALVNVILKQANEDQEEAWRQIDEAFLE
Dmel_Dsx        399 RVEI--------------------------------------------------NRTVAQ
Lcup_Dsx        368 IQVLKQY----NLNIYDGNELRK---------------LKTERRYENHLARSECDETIKQ
Tcas_Dsx        199 RDT--EI----LLDFCQRLKDK---FQLSWKMISLVDVILKYA-KDQDEAWRQIDEAFLE
Bmor_Dsx        244 KMIVDEYARKHNLNVFDGLELRNSTRQ---------------------------------
Ekue_Dsx        214 HWMVHQW----RLSLCSLLQAR---KE---------------------------------
Tdom_Dsx        238 QAEL--------------------------------------------------------
Dmel_Dmrt99B    418 EDL---------------------------------------GQTPPQVP-----PSPP-
Lcup_Dmrt99B    385 EDIL-----------------------------------TNSTNSPPNVP-----PSPP-
Tcas_Dmrt99B    296 SHE---------------------------------------------------DATPP-
Bmor_Dmrt99B    272 PEP-----------------------------------------NPYHVT----QESPPY
Bmut_Dmrt1      261 -----------------------------------------ENR---HAV------SSQ-
Drer_Dmrt1      183 -------------------------------------------RLSSHNV------SPQ-
Mmus_Dmrt1          ------------------------------------------------------------
Xlae_Dmrt1      235 -----------------------------------------ENRFPGHSG------SSQ-
```

```
                  721 .......730.......740.......750.......760.......770.......780
Haxy_Dsx          243 VRTWAAVEAARSA-----------YRHIPYAGFYSTATALYHPHMYLPAIPTYHS-----
Dmel_Dsx          409 I------------------Y----YNYYTPMAL------VNGAPMYL-TYPSIEQGRY--
Lcup_Dsx          409 IR---LKEATEQLNQLTQTY----YNYQRYGTL---------PPAYW-AYPSIQLGRTIW
Tcas_Dsx          249 IRALAAVEAARYT-----------YHHIPYSGLYPNAATAIYPPVYLPSMSMYH------
Bmor_Dsx          271 ------------------------YGL---------------------------------
Ekue_Dsx          234 ------------------------YSMSC-----CSPRFVIAPE-YA-------------
Tdom_Dsx          242 -RSMALREAARVIHYPGPYY----YNYYP-------------PTPYLPTPPSIDL-----
Dmel_Dmrt99B      433 ---FPMKSAFSPLVPP-SVFGSPTHRYHPFMQA-HAKRFLTAP--YA-------------
Lcup_Dmrt99B      404 ---FPLKSAFSPLVPPAAVFGSPTHRYPPFMQA-HAKRFLTAP--YA-------------
Tcas_Dmrt99B      304 -------SAFSPLGPP-TNF----HRFSP------SRRFLSAP--YA-------------
Bmor_Dmrt99B      287 LQNYQSKSAFSPLSNQ-------SFKFSP------SRRFLTPP--YS-------------
Bmut_Dmrt1        270 ---YRM------------------HSYYP-------------PPSYL-------------
Drer_Dmrt1        193 ---YRT------------------HSYYS-------------S--YL-------------
Mmus_Dmrt1            ------------------------------------------------------------
Xlae_Dmrt1        247 ---FRM------------------HSYYP-------------P--YL-------------

                  781 .......790.......800.......810.......820.......830.......840
Haxy_Dsx          287 -----------------GADLLPTVPSRS-PPLPQAPSLTPPQSN--AVLRPGSRA----
Dmel_Dsx          438 ----GAHFTHLPLTQICPPTPEPLALSRS----PSSPSGPSAVHNQ-KPSRPGSSNG---
Lcup_Dsx          452 TELPTPHFAAIIPPHSAPTPEEPLTLSRA----STSPS---------KISRSGSSSICGE
Tcas_Dsx          292 -----------------PATLLGSVPTST------SPSHSPPIVP--RAIRPSSRA----
Bmor_Dsx              ------------------------------------------------------------
Ekue_Dsx          251 -----------------PLLPLPLTTQRP----SPPPAHL--------------------
Tdom_Dsx          279 -----------------PSYPPPLIPHVS-TGLAAAPE----------------------
Dmel_Dmrt99B      473 -----------------GTGYLPGVLSAA------------------DIEQSESNG----
Lcup_Dmrt99B      445 -----------------GTGYLPGVINPA------------------DVEQSESN-----
Tcas_Dmrt99B      331 -----------------GTGYLPTVIRPP-------PDYLSMVGSVHDIYSSDKT-----
Bmor_Dmrt99B      319 -----------------GTGYLPTVIRPP-------PEYLSSFMPPSELMYGRQL-----
Bmut_Dmrt1        283 -----------------GQSMSQ------IFTFEDSASYSEAKASVFSPPSSQDS-----
Drer_Dmrt1        204 -----------------SQGLGA-ACVQP-------STCPEPKAAAFS-DGAQDS-----
Mmus_Dmrt1            ------------------------------------------------------------
Xlae_Dmrt1        258 -----------------GQSVPNPACVPPFLTFEEIPSYSEAKASVLSPPSSQDS-----

                  841 .......850.......860.......870.......880.......890.......900
Haxy_Dsx              ------------------------------------------------------------
Dmel_Dsx          486 TVHSAASPTMVTTMATTSSTPTLSRRQRSRSATPTTPPPPPPAHSSSNGAYHHGHHLVSS
Lcup_Dsx          499 SITATSTPTPTTT--TTPSAGVIAAAAAAAAAAAAT------------------------
Tcas_Dsx              ------------------------------------------------------------
Bmor_Dsx              ------------------------------------------------------------
Ekue_Dsx              ------------------------------------------------------------
Tdom_Dsx          299 ------SPRLIDGRAHTNTSTSAS------------------------------------
Dmel_Dmrt99B      494 --AGGIGLDRTSNAGDSQD-----------------------------------------
Lcup_Dmrt99B      465 ---GGASVDRNSNAGDSQD-----------------------------------------
Tcas_Dmrt99B      362 ---SASSPGSNTSSDKTSYSE---------------------------------------
Bmor_Dmrt99B      350 ---QVPSPGTSPTSDNTNNDGFSD------------------------------------
Bmut_Dmrt1        315 ---GLVSLPSSSPIGNESTKAVLDCESASEPSNFAVAP-------------------IIE
Drer_Dmrt1        233 -----VSISSMINAENK-----LECESSSESGSFSVDS-------------------IIE
Mmus_Dmrt1            ------------------------------------------------------------
Xlae_Dmrt1        296 ---GVISLSSNSPVSNESTKAVAEQEPNSESSLFTVTT-------------------AAE

                  901 ....
Haxy_Dsx              ----
Dmel_Dsx          546 TAAT
Lcup_Dsx              ----
Tcas_Dsx              ----
Bmor_Dsx              ----
Ekue_Dsx              ----
Tdom_Dsx              ----
Dmel_Dmrt99B          ----
Lcup_Dmrt99B          ----
Tcas_Dmrt99B          ----
Bmor_Dmrt99B          ----
Bmut_Dmrt1        353 EDE-
Drer_Dmrt1        264 GATK
Mmus_Dmrt1            ----
Xlae_Dmrt1        334 NGE-
```
