## Supplementary figures and images for "Sex-specific *doublesex* regulation targeting the color-patterning gene *h* underlies the evolution of wing sexual dimorphism in the harlequin ladybug *Harmonia axyridis*"

### Supplementary Figure 2

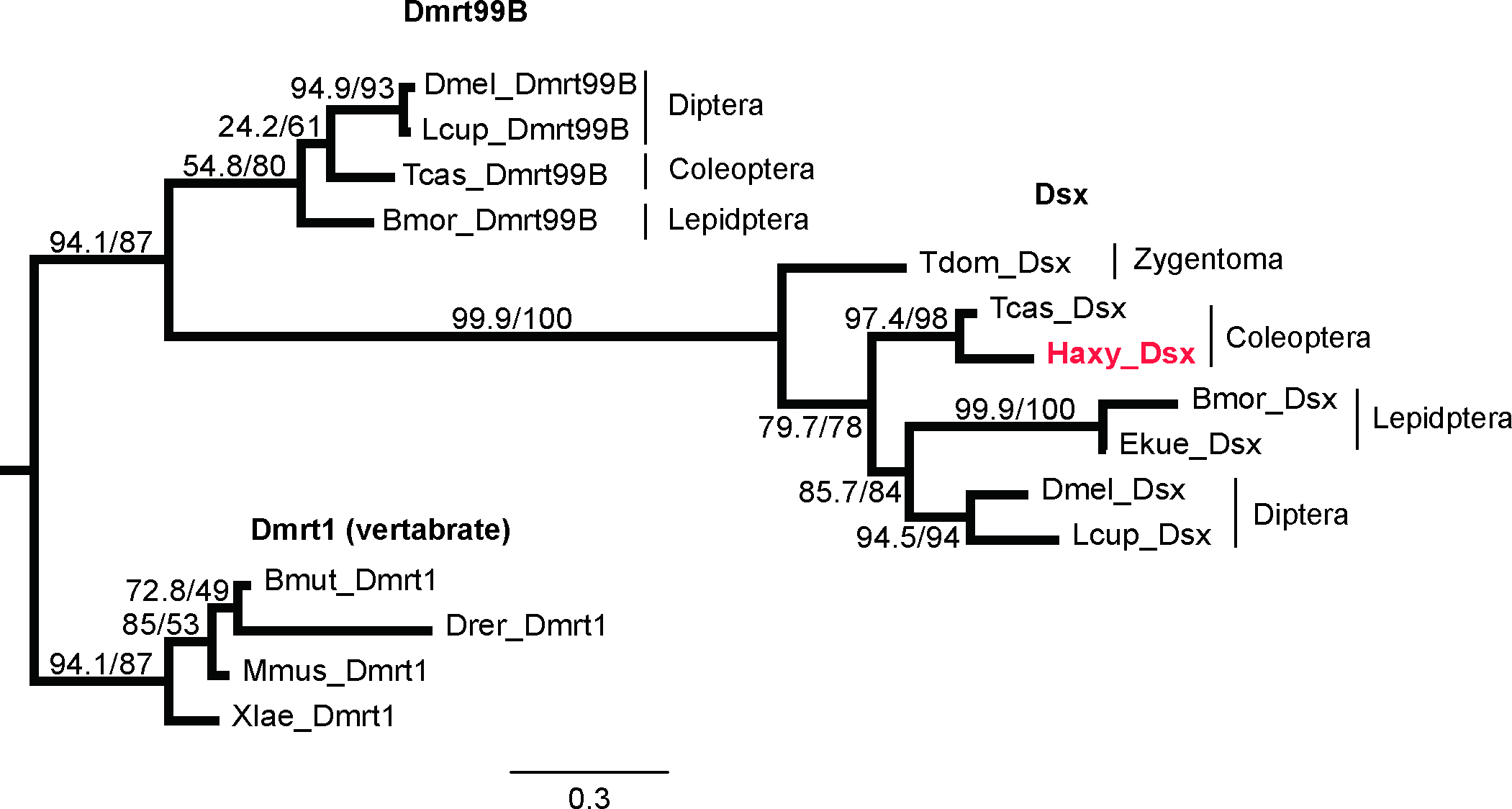

### Supplementary Figure 3

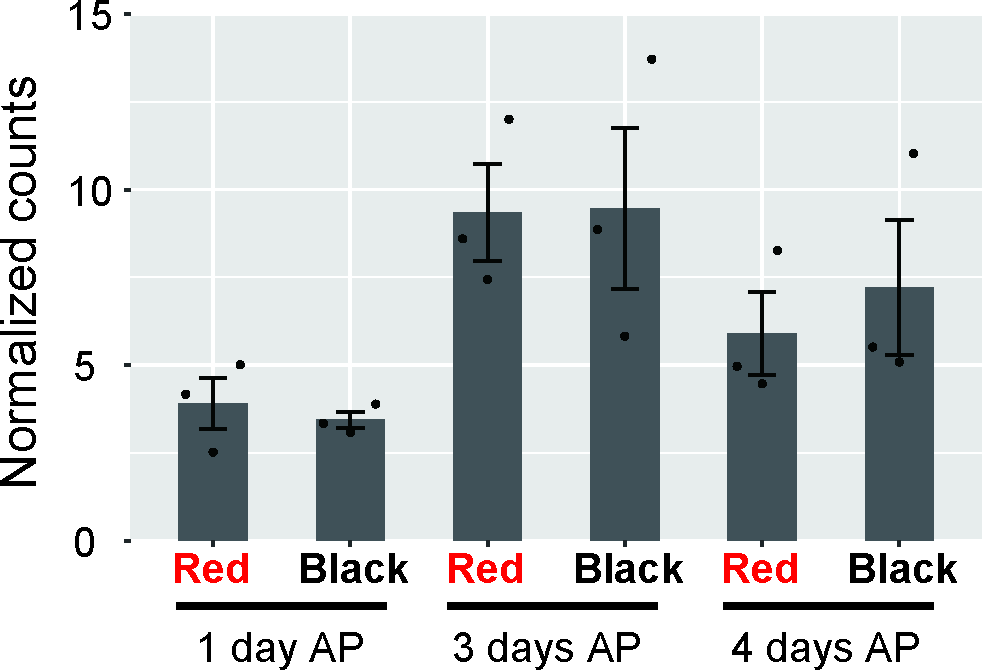

### Supplementary Figure 4

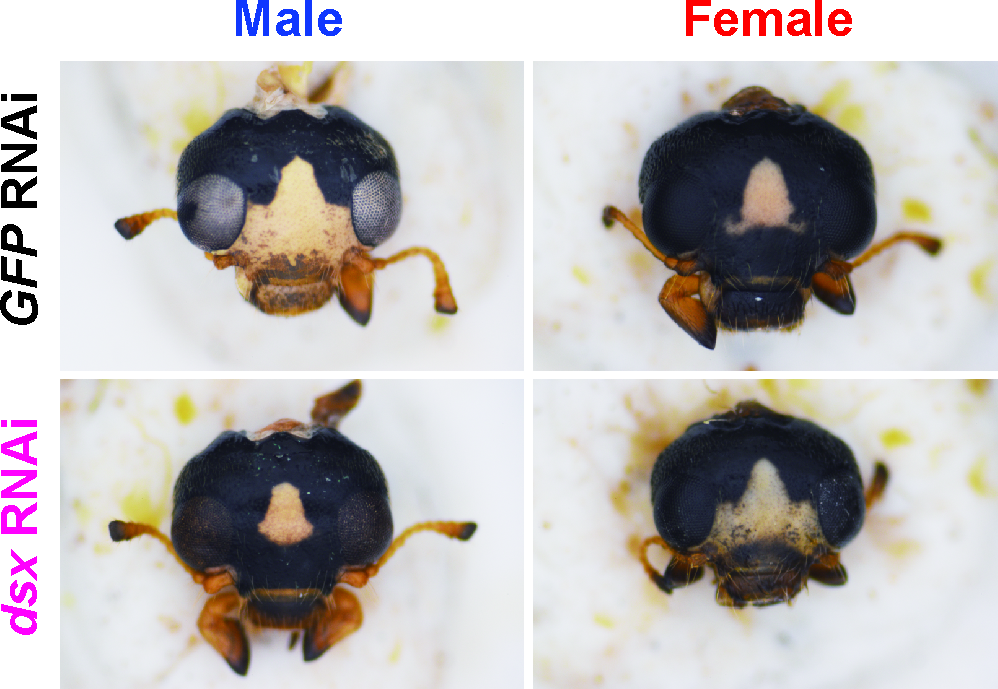

### Supplementary Figure 5

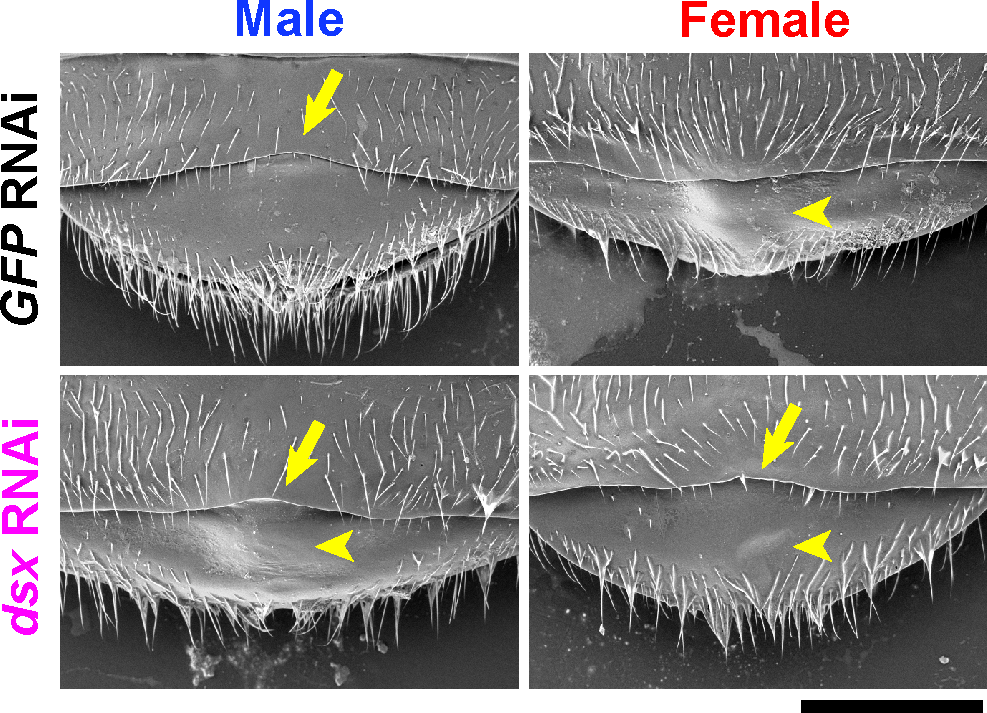

### Supplementary Figure 6

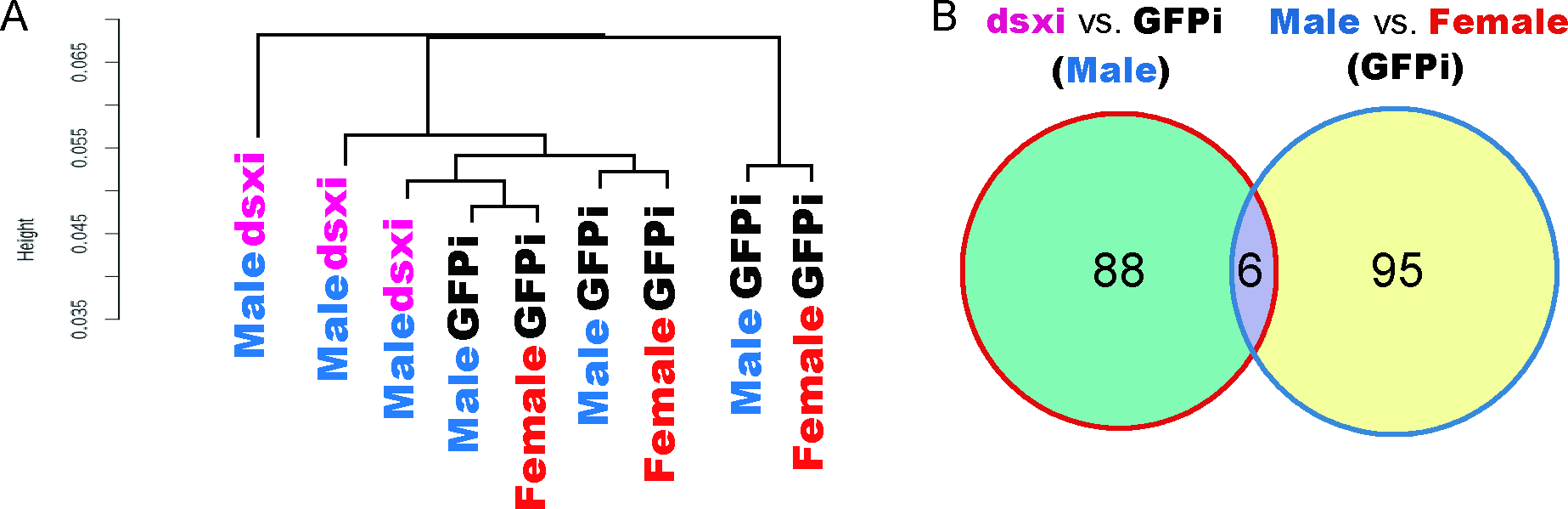

### Supplementary Figure 7

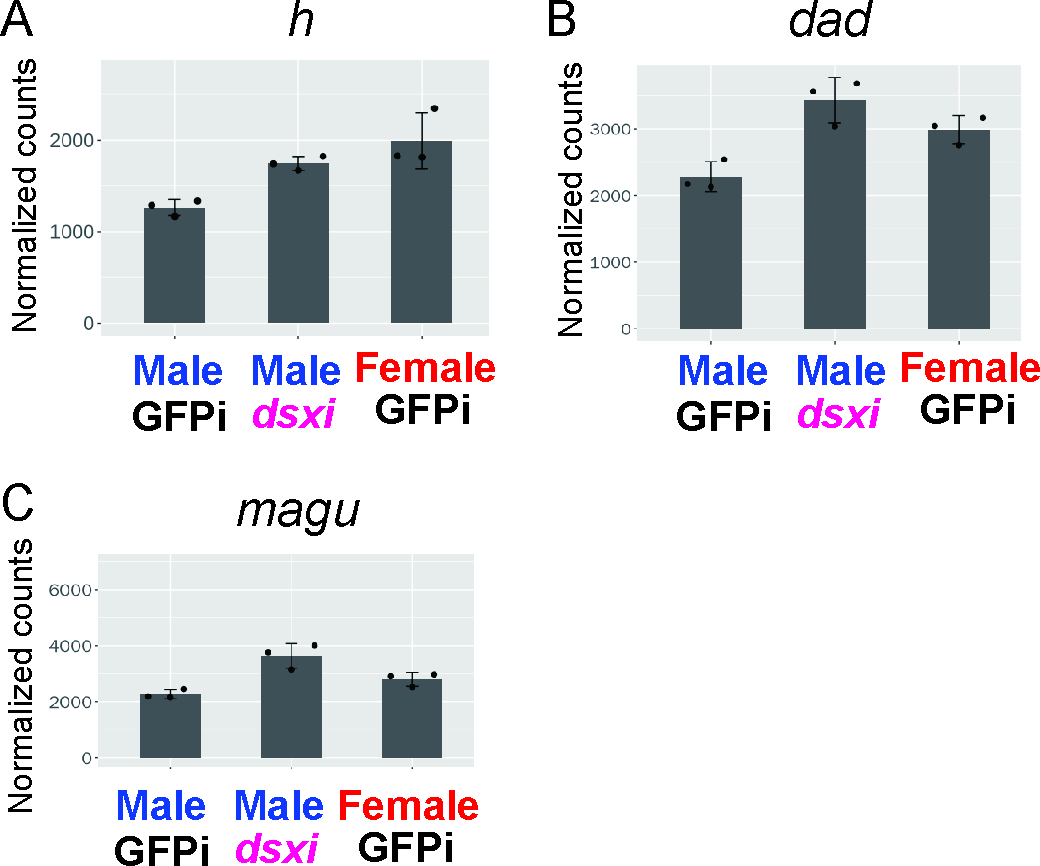

### Supplementary Figure 8

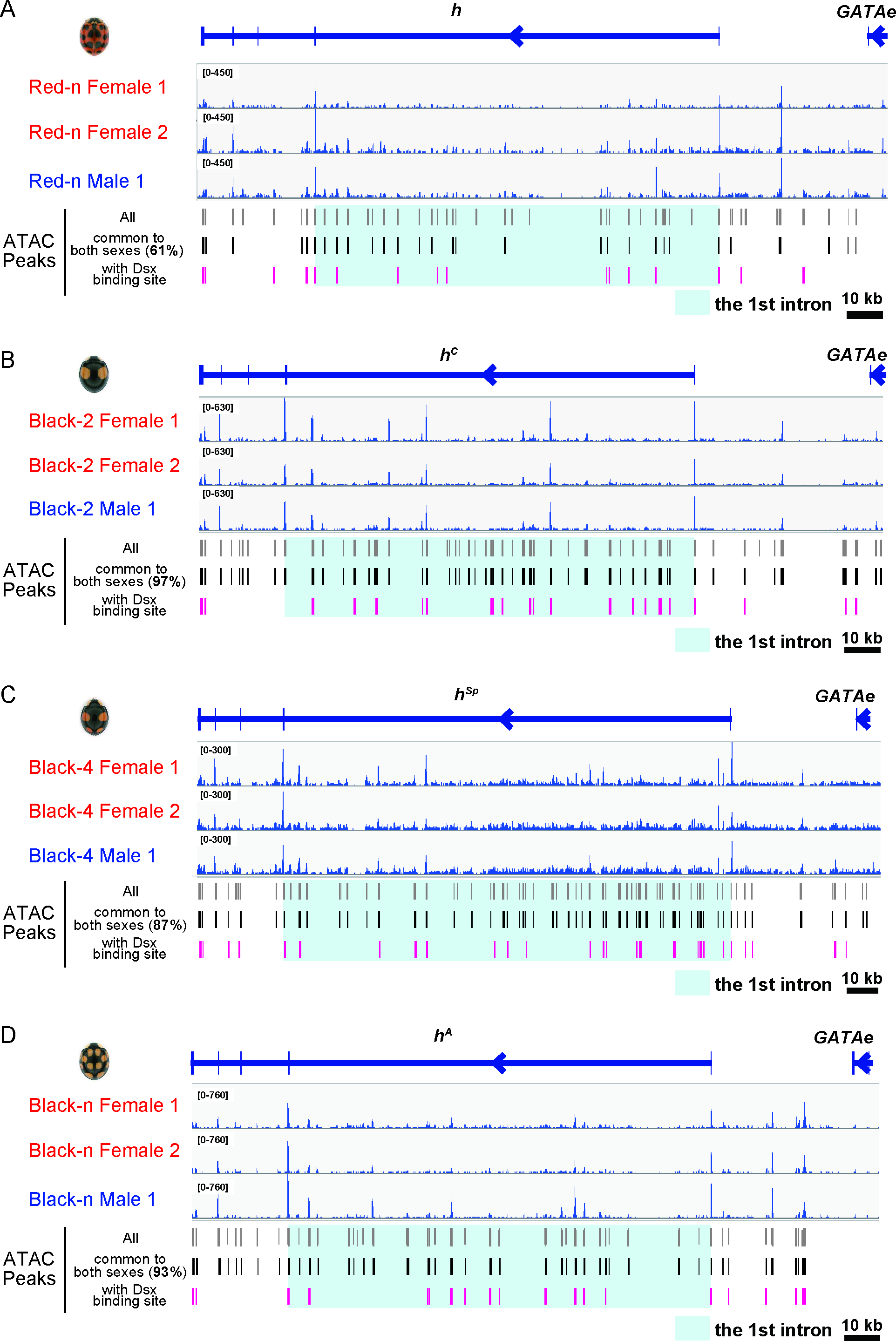
